# On Human Motor Coordination: The Synergy Expansion Hypothesis

**DOI:** 10.1101/2024.04.10.588877

**Authors:** F. Tessari, A.M. West, N. Hogan

**Affiliations:** Department of Mechanical Engineering, Massachusetts Institute of Technology, Cambridge, 02139, USA; Department of Brain and Cognitive Sciences, Massachusetts Institute of Technology, Cambridge, 02139, USA

## Abstract

The search for an answer to Bernstein’s degrees of freedom problem has propelled a large portion of research studies in human motor control over the past six decades. Different theories have been developed to explain how humans might use their incredibly complex neuro-musculo-skeletal system with astonishing ease. Among these theories, motor synergies appeared as one possible explanation. In this work, the authors investigate the nature and role of synergies and propose a new theoretical framework, namely the “expansion hypothesis”, to answer Bernstein’s problem. The expansion hypothesis is articulated in three propositions: mechanical, developmental, and behavioral. Each proposition addresses a different question on the nature of synergies: (i) How many synergies can humans have? (ii) How do we learn and develop synergies? (iii) How do we use synergies? An example numerical simulation is presented and analyzed to clarify the hypothesis propositions. The expansion hypothesis is contextualized with respect to the existing literature on motor synergies both in healthy and impaired individuals, as well as other prominent theories in human motor control and development. The expansion hypothesis provides a novel framework to better comprehend and explain the nature, use and evolution of human motor skills.

**Significance Statement:** Understanding how humans effortlessly control coordinated movements has been a long-standing challenge in neuroscience. This research introduces the “expansion hypothesis”, a new framework to explain how we develop, learn, and use motor synergies – coordinated activities of multiple features such as joints and muscles – that simplify movement control. By breaking down the nature of these synergies into mechanical, developmental, and behavioral aspects, this study offers novel insights into how our brains and bodies work together to achieve fluid motion. This work not only advances the scientific understanding of human motor control but also has potential implications for improving rehabilitation strategies for individuals with movement impairments and for developing more dexterous human-inspired robotic control techniques.

## Introduction

In 1899, Prof. Robert Woodworth questioned how humans could coordinate the actions of their incredibly numerous muscles and joints to perform most of their motor tasks with astonishing ease. Citing his own words: *“Even in comparatively unskilled movements it is remarkable how many groups of muscles must cooperate, and with what accuracy each must do just so much and no more*.*”* (Woodworth, 1899).

Similarly, more than half a century later, Prof. Nikolai Bernstein published another milestone contribution trying to answer the same research question. Interestingly, his research started with a similar opening: “*The science of human movements and theories of motor co-ordination have until recently been one of the most obscure branches of human and comparative physiology;* …” (Bernstein, 1967). Bernstein later formulated what is known as the “degrees of freedom” problem. This concerns the astonishing redundancy (or abundance) of the human motor control system that fundamentally affords infinitely many solutions to the same motor problem. Therefore, the natural question that arises is: how do humans compute, find or select their specific solution for a given motor task?

Multiple hypotheses have been proposed through the years to answer this question. These hypotheses can be classified into four main categories – not always mutually exclusive: optimal control hypotheses (Kumar et al., 2016; Scott, 2004); equilibrium point hypotheses (Feldman, 1966, 1986); uncontrolled manifold hypotheses (Kang et al., 2004; Scholz & Schöner, 1999; Todorov, 2004); and motor synergy hypotheses (D’Avella & Bizzi, 2005; Latash et al., 2007; Santello et al., 1998).

Here we focus our attention on the theory of motor synergies. The term “synergy” – from the Greek word συνεργία i.e., ‘working together’ – was initially adopted to describe particularly repetitive and coordinated motor tasks such as locomotion. However, in more recent years the term has found an increasing use in the motor control and medical literature in a broader sense and – often times – with different connotations.

Depending on the nature of the analyzed human behavior, we can talk of either neural (Cheung & Seki, 2021; Santello et al., 2013), muscular (Cheung et al., 2009; D’Avella & Bizzi, 2005; D’avella & Tresch, 2001; Weiss & Flanders, 2004), or kinematic synergies (Santello et al., 1998, 2016; Todorov & Ghahramani, 2004). In all these contexts, synergies are interpreted as a dimensionality reduction method that humans adopt to simplify control by fundamentally transforming a high-dimensional task to a lower-dimensional one. A common example can be found in the many studies of the kinematics of the human hand (Mason et al., 2001; Santello et al., 1998, 2013, 2016; Todorov & Ghahramani, 2004; Weiss & Flanders, 2004; West et al., 2023). By extracting synergies through dimensionality reduction techniques (e.g., Principal Component Analysis), a few synergies were able to account for most of the variance of different reaching and grasping hand motions. In a similar fashion, synergies were also extracted in studies of posture and movement of the lower limbs at the muscular level (Ting & McKay, 2007; Yang et al., 2019). An important question, rarely addressed in this prior work, is how these synergies emerge. This paper explores a possible answer.

Moreover, in all the aforementioned motor control studies, synergies assume a mostly-positive connotation. They represent a way to reduce control complexity by voluntarily reducing the dimensionality of the analyzed motor tasks. However, it is important to mention that synergies can also assume a negative connotation, such as in neuro-rehabilitation. In these cases, synergies or synergistic actions are interpreted as the unwanted, and typically uncontrolled, movements of patients’ limbs in predetermined coordination patterns. For example, when discussing stroke survivors, synergy is often used to describe the result of undesired and involuntary motion of the paretic side (Avni et al., 2022; Barroso et al., 2017; Dipietro et al., 2009; Hadjiosif et al., 2022; Krakauer & Carmichael, 2017; Neckel et al., 2006; Roh et al., 2013; Yang et al., 2019).

This apparent contrast of positive and negative connotations of synergies often leads to confusion and disagreement between motor control and rehabilitation experts. In this work, to minimize confusion, we start by proposing a definition of motor synergy i.e., *the coordinated and simultaneous activity of a finite group of features (neural, muscular, kinematic) to perform a specific motor task*. This definition is intended to remove any value judgement from the concept of synergy, thus leaving the reader free to consider it as negative or positive depending on the application context.

The current mathematical methods to extract synergies are based on matrix factorization techniques such as Principal Component Analysis (PCA), Factor Analysis (FA), Non-negative Matrix Factorization (NNMF), or Singular Value Decomposition (SVD) (D’avella & Tresch, 2001; Fazle Rabbi et al., 2020; Todorov & Ghahramani, 2004; Tresch et al., 2006; West et al., 2023). All of these methods are fundamentally designed to find lower-dimensional spaces within a given hyper-space of analyzed features. For example, when studying human hand kinematics, the feature space might account for ∼20 hand joint angles. In this 20-dimensional hyper-space, the aforementioned techniques numerically extract directions (synergies) where most of the variance of the feature space lies.

These approaches intrinsically limit the number of possible synergies to no more than the number of analyzed features. However, this intrinsic limitation – dictated by the adopted mathematical approaches – precludes an answer to several interesting questions: How do synergies emerge in human behavior? How many synergies can humans have? Is there an upper or lower limit? The former question is sparsely investigated. If a synergy is a coordinated action of multiple features, how do we reach this coordination? Do we start from individual joint or muscle actions and learn to combine them? Or do we move from highly coordinated joint or muscle patterns and learn to refine them?

In this work, the authors propose a “synergy expansion hypothesis”, to answer Bernstein’s problem. This hypothesis aims to provide a comprehensive answer to the aforementioned questions by grounding the motor synergy hypothesis in the biomechanical structure and developmental history of human motor behavior. A numerical example is provided to facilitate an intuitive understanding of the hypothesis.

### The Synergy Expansion Hypothesis

Before introducing the propositions of the synergy expansion hypothesis, it is important to provide readers with a common framework and related nomenclature to contextualize/locate synergies in the literature of motor planning and execution. We can define a ‘*motor task*’ as any task requiring the temporal coordination of multiple features (muscles or joints) of the human musculoskeletal system. For example, a motor task could be a reach-and-grasp movement or the movement performed when swinging a golf club, throwing a football or playing a musical instrument. Such a motor task can be decomposed as a ‘*sequence*’ of temporally grouped ‘*motor chunks*. For a review of the theory of motor sequencing and chunking, we invite readers to refer to the works of Lashley (original) and Diedrichsen and Kornysheva (more recent) (Diedrichsen & Kornysheva, 2015; Lashley, 1951). In the context of this work, a ‘*motor chunk*’ can be imagined as an intermediate step of decomposition of a ‘*motor task*’ that can be represented by a continuous temporal evolution ***c***_***j***_(*t*). For example, in piano playing (or a finger tapping sequence), a motor chunk can represent a sub-sequence of the finger movements that has its own distinct temporal pattern (Diedrichsen & Kornysheva, 2015; Wiestler & Diedrichsen, 2013; Yokoi & Diedrichsen, 2019).

Each ‘*motor chunk*’ can in turn be decomposed into a series of ‘*elementary actions’* representing the fundamental ‘atomic’ elements composing motion. These ‘*elementary actions*’ are made of two distinct parts: the ‘*synergy composition*’ vector ***w***_*i*_ (which contains the information of the selection of each feature – muscle or joint – to the given motor action), and the ‘*temporal evolution*’ *u*_*i*_(*t* − *t*_*i*_) (which represents the time trajectory of the synergy composition vector). As a result, a generic ‘*motor task’* that runs from an initial time 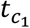 to a final time *t*_*f*_ can be formalized as:

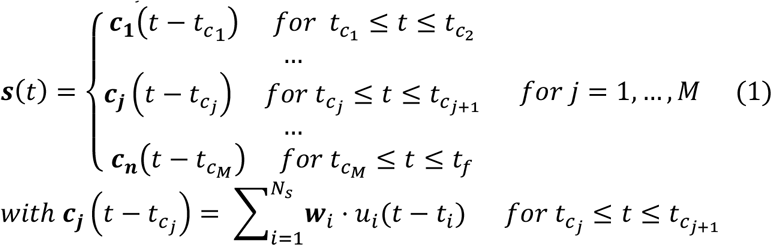

where ***s***(*t*) represents the ‘motor task’ sequence composed by *M motor chunks* 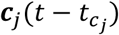, which are in turn the result of the summation of the *N*_*s*_ *synergy compositions* ***w***_*i*_ each multiplied by their own *temporal evolution u*_*i*_(*t* − *t*_*i*_). Figure 1 provides a schematic representation of the proposed hierarchical organization for the planning and execution of a given motor task.

**Figure 1.**
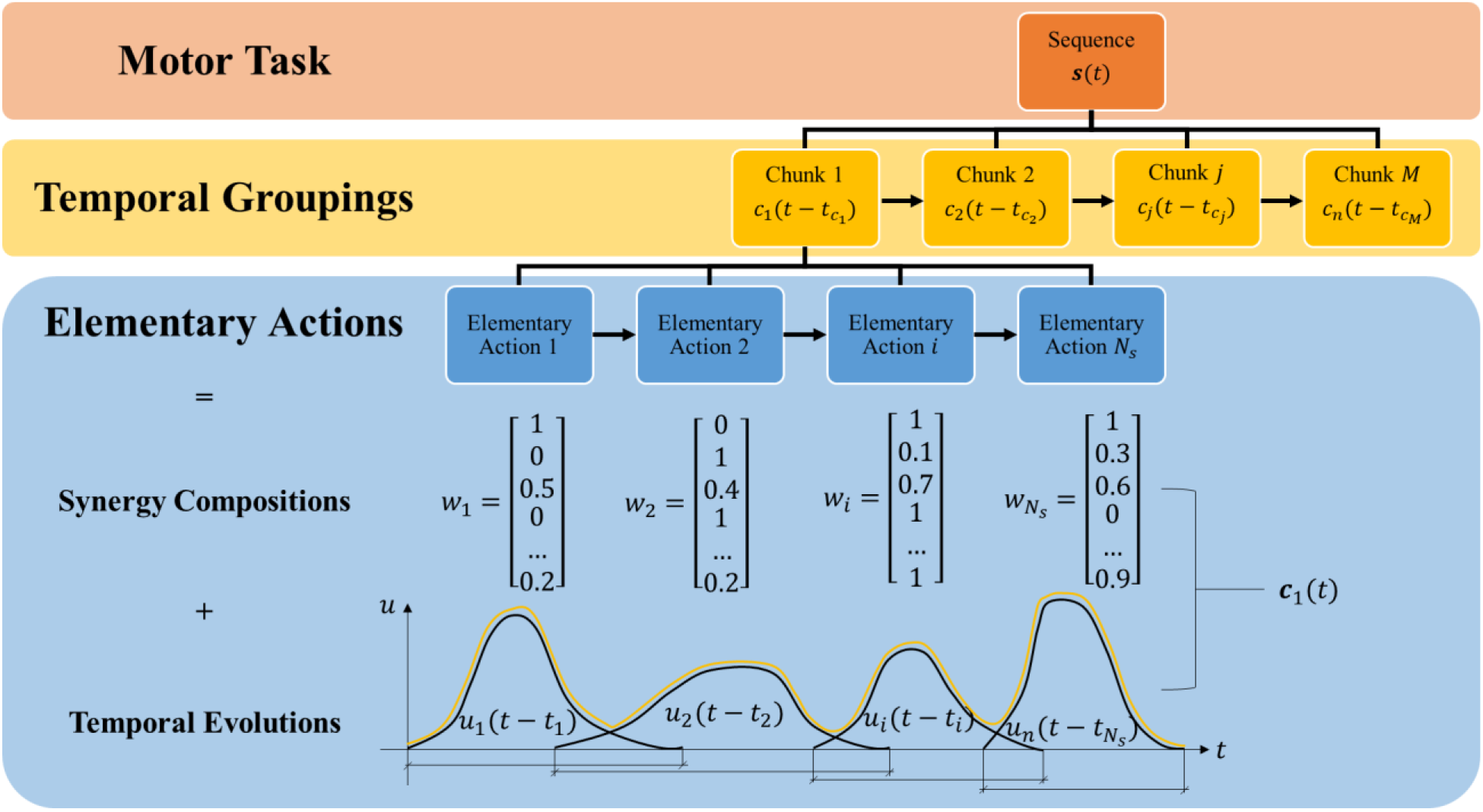
Hierarchical organization of motor planning and execution. Each motor task can be represented by a sequence (***s***(*t*)) of motor chunks. These motor chunks 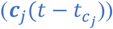 represent temporally grouped elementary actions. Each elementary action can be decomposed into a synergy composition vector ***w***_*i*_ – containing the relative contribution of each feature ‘*f’* (muscle or joint) to the elementary action – and a temporal evolution trajectory *u*_*i*_(*t* − *t*_*i*_) – representing the temporal pattern of the associated synergy composition vector.

Such a hierarchical organization of motor planning and execution has been widely recognized in many studies of the human neuromotor control system (Loeb et al., 1999; Merel et al., 2019; Yokoi & Diedrichsen, 2019).

It is worth emphasizing that the proposed hierarchical framework assumes that the synergy composition is a time-invariant, finite length, vector. This is in line with the work of Hart and Giszter who experimentally showed how synergies are better represented by time-invariant quantities (Hart & Giszter, 2013). Moreover, it is also worth noting that the temporal evolutions of the elementary actions (*u*_*i*_(*t* − *t*_*i*_)) can present temporal overlap. In other words, it is theoretically possible to command a first elementary action, and then fire a subsequent one without waiting for the first one to be completed. This is in agreement with frequently observed pre-reaching adaptations of the hand configuration while performing reaching motions as well as by the phenomenon of blending of submovements in stroke recovery (Dipietro et al., 2009; Flash & Henis, 1991; Rohrer et al., 2004).

In light of the presented hierarchical framework, the synergy expansion hypothesis can be articulated as three propositions:

1. **Mechanical**: The number of synergies is not limited by the number of controlled features.
2. **Developmental**: Humans start from highly-stereotyped crude coordination patterns (primordial synergies) and, through motor learning and development, expand to new and more specialized synergies.
3. **Behavioral**: Learned synergies are evoked upon need during a given elementary action.

### Mechanical Proposition

The human motor control system can be described in at least three major geometric hyper-spaces: task space, joint space and muscle space (Figure 2). The task (or work) space is the physical space in which we perform our motor actions. It can be described, considering a single rigid end-effector, by up to *n*_*ts*_ = 6 independent variables. The most common (but not unique) choice is an orthogonal reference frame of three translations (surge, sway, heave i.e., *x, y, z*) and related axial rotations (roll, pitch, and yaw i.e., *α, β, γ*). These variables can be associated with a given “flow” i.e., task-space velocities (***v***), and a given “effort” i.e., task-space generalized forces (***F***) (Lagrange, 1788; Newton, 1687; Paynter, 1961). The joint (or configuration) space is the hyper-space defined by the human articulations/joints or, in other words, skeletal degrees of freedom. This hyper-space has as many dimensions as the number of human degrees of freedom. That number varies depending on the definition of “joint”, ranging from *n*_*js*_ ≈ 200 (Kandel et al., 2021) to about *n*_*js*_ ≈ 350 (Cleveland Clinic, 2024; Healthline, 2024). The strictest criterion, for example, considers any two consecutive bones as a “joint”. In the joint space, we can define a generalized coordinate (*q*) for each degree of freedom and this typically (but not exclusively) coincides with the joint angular configuration. Similarly to task-space, we can define a vector of joint-space velocities 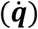 and of joint-space generalized forces (***τ***) representing the “flow” and “effort” variables in the joint space (Lagrange, 1788; Newton, 1687; Paynter, 1961). Finally, we have the muscle hyper-space. This comprises a number of dimensions equal to the number of human muscles. This number is also subject to variation depending on the selection criterion, but it can be reasonably stated that there are *n*_*ms*_ ≈ 600+ independently-addressable muscles in the human body (Kandel et al., 2021). The muscle hyper-space can be, therefore, characterized by a vector of muscle displacements (***x***_***m***_), muscle velocities 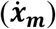 and muscle forces (***F***_***m***_).

**Figure 2.**
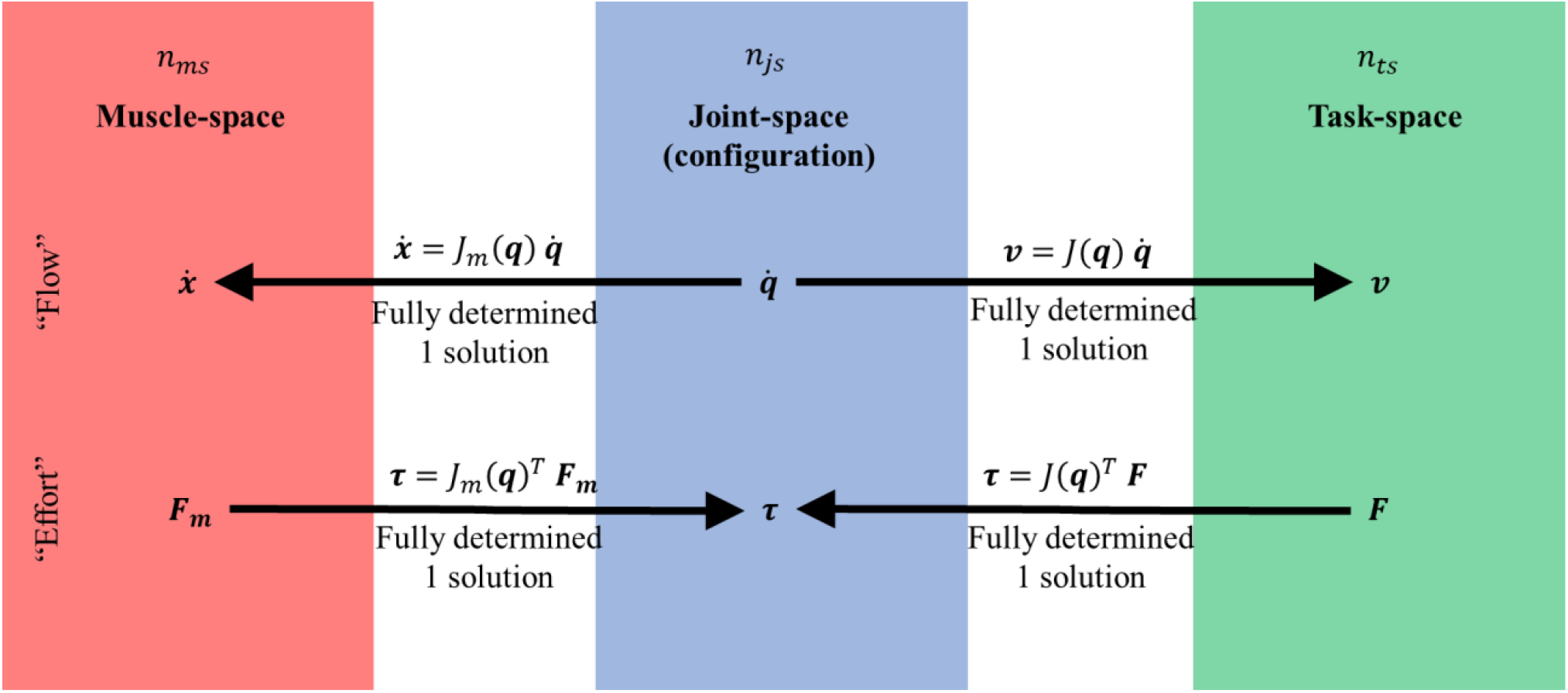
The relationship between some of the different spaces of human motor control. These spaces are characterized by *n*_*ms*_, *n*_*js*_, *n*_*ts*_ dimensions respectively for the muscle, joint and task space. It is important to underline that *n*_*ms*_ > *n*_*js*_ > *n*_*ts*_. The arrows describe the direction of fully determined transformations. Interestingly, the joint-space coordinates (*q*) allow for a full description of both the muscle and task space variables.

Knowing the skeletal and muscular geometry of the human body, it is possible to define a mapping between the effort and flow variables of the task, joint, and muscle spaces. Specifically, the joint space coordinates (***q***) allow the unique definition of both the task and muscle space coordinates. The differential form of these two mappings is provided by two different Jacobian matrices: the task-space Jacobian (*J*(***q***)) and the muscle-space Jacobian (*J*_*m*_(***q***)). For a detailed presentation of Jacobian matrices, the authors recommend the work by Murray et al. (Murray et al., 1994).

The task-space Jacobian is a *n*_*ts*_ × *n*_*js*_ matrix 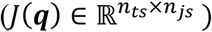 that uniquely maps the joint-space velocities 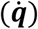 to task-space (***v***):

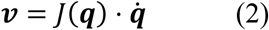

The muscle-space Jacobian is a *n*_*ms*_ × *n*_*js*_ matrix 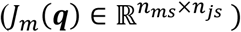 that uniquely maps the joint-space velocities 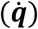 to muscle-space 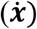:

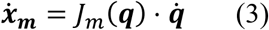

These mappings focus on the “flow” variables of the three considered spaces. However, the three spaces are also physically coupled and, as consequence, obey the law of power continuity. The mechanical power at the task-space (*P*_*t*_ = ***F*** · ***v***), joint-space 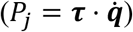, and muscle-space 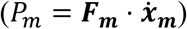 must be the same:

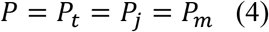

Substituting Eqs. 2 and 3 into Eq. 4, it is possible to obtain the mapping between the “effort” variables of the considered hyper-spaces:

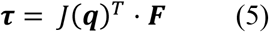

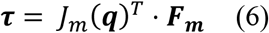

As can be observed from Eqs. 5-6, the transpose of the task and muscle Jacobians provides a unique mapping between the task-space and muscle-space forces to the joint torques. Figure 2 shows a graphical representation of these relations.

These unique mappings fundamentally provide a one-to-one relation between the different hyper-spaces of human motor action. However, such mappings are non-consecutive i.e., you can’t directly map from task-space to muscle-space – either in terms of “flow” or “effort” variables – with a unique transformation.

For example, if we try to identify the muscle velocities 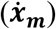 required to perform a desired task-space motion (***v***), we need to combine Eqs. 2 and 3, obtaining:

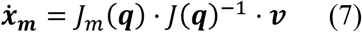

This requires the matrix inversion of the task-space Jacobian. However, because *n*_*js*_ > *n*_*ts*_, the resulting system of equations is under-determined, and therefore, there is no unique inverse, i.e., solution, but infinitely many of them. Specifically, there exist infinitely many solutions whose dimensionality is equal to the size of the null-space of the task-space Jacobian (*Null*(*J*(***q***))), where the null-space is the linear subspace that describes a mapping to a zero vector.

Similarly, if we try to identify the muscle force vector (***F***_***m***_) required to generate a desired task-space force (***F***), we need to combine Eqs. 5 and 6, obtaining:

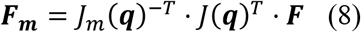

In this case, the inversion of the muscle-space Jacobian transpose also leads to an under-determined system of equations with infinitely many solutions (with dimensionality determined by the null-space of the muscle-space Jacobian *Null*(*J*_*m*_(***q***))) since *n*_*ms*_ > *n*_*js*_.

From a human motor control perspective, we can imagine these non-unique solutions as the spaces where kinematic and muscular synergies may usefully be defined. Independently of the neural basis of the synergies, to generate a desired task-space force or motion, a specific trajectory in the null-space must be selected in agreement with Equations 7 and 8. The selected trajectories in the null-space represent our selectable synergies. In fact, a synergy can be imagined as one of the possible solutions of the Jacobian inverse in the “flow” domain or transposed Jacobian inverse in the “effort” domain. Specifically, kinematic synergies can be identified in the inversion of the task-space Jacobian, *J*(***q***), when going from task-space velocities to joint-space velocities 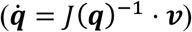, while muscular synergies may be identified when mapping joint-space torques to muscle-space forces (***F***_***m***_ = *J*_*m*_(***q***)^−*T*^ · ***τ***). These coordination patterns can be considered as **selectable synergies** in the sense that individuals can choose between (infinitely) many of them and select the one they prefer. Figure 3 shows a graphical description of the emergence of selectable kinematic and muscular synergies from the matrix inversions of the Jacobians.

**Figure 3.**
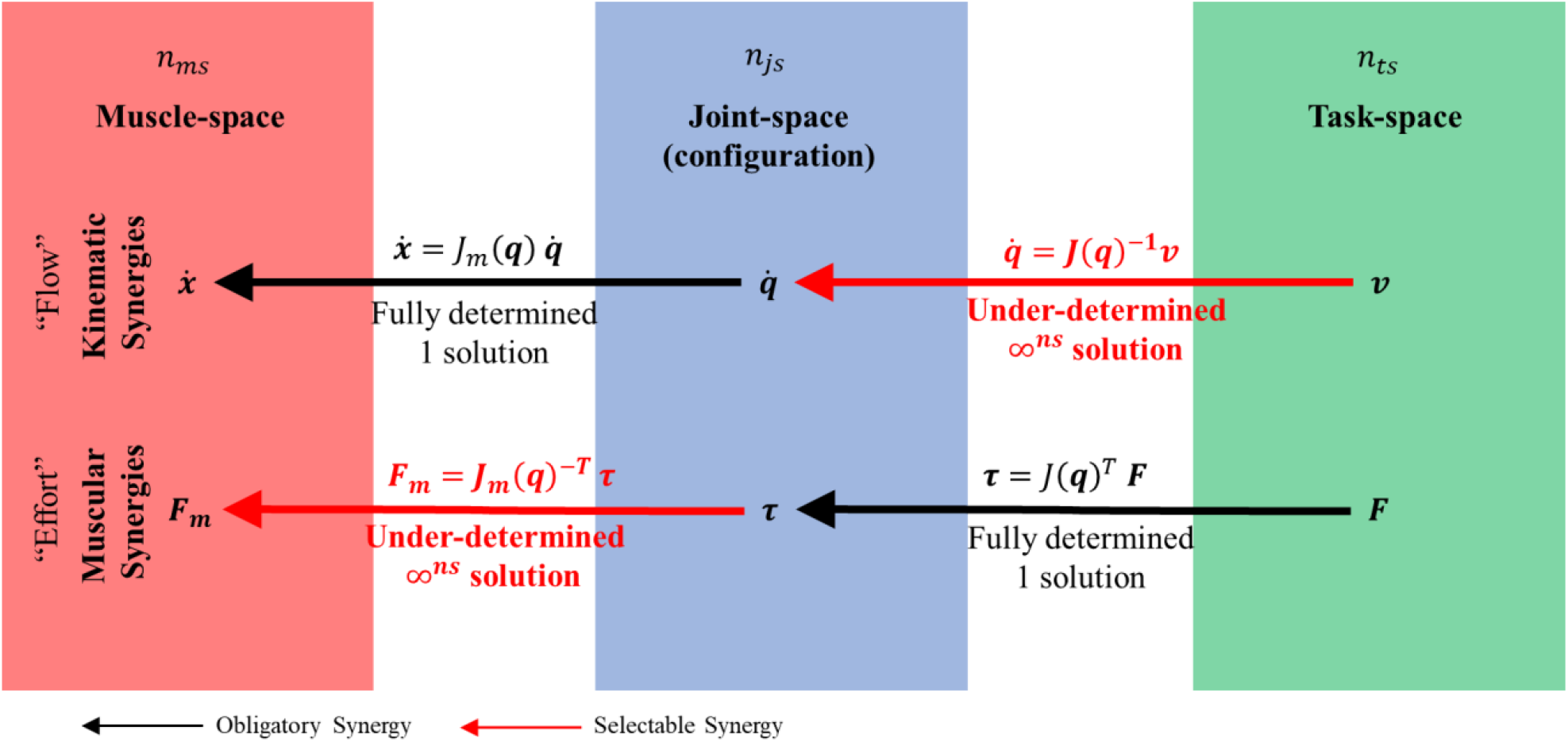
The mechanical framework describing the emergence of motor control synergies. Synergies emerge from the mapping of task-space variables into joint, and subsequently, muscle space quantities. Synergies can be categorized as obligatory or selectable, depending on the determination of the related transformation i.e., a fully-determined transformation will lead to an **obligatory** synergy, while an under-determined transformation will lead to a **selectable** synergy.

It is interesting to observe that, when moving from task-space to muscle space, there are still some uniquely defined mappings (Fig. 3). For example, joint-space torques are uniquely defined by task-space forces (***τ*** = *J*(***q***) · ***F***); muscle-space velocities are uniquely defined by joint-space velocities 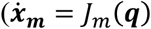. These uniquely-defined transformations are dictated by geometry and configuration, and can be interpreted as **obligatory or mandatory synergies**. In simpler words, if I want – for example – to generate a certain end-effector force with my arm given a certain configuration, I will have to generate a uniquely defined set of joint torques with no option for other solutions.

A limitation – worth emphasizing – of the mechanical proposition is that the proposed framework in Figure 3 does not consider that the lever arms of the muscle forces are not always constant, and are functions of the exerted force. While this would influence the relation between force and torque, it does not change the fact that an under-determined mapping exists between muscle forces and joint torques. Therefore, considerations of the first proposition of the expansion hypothesis are still valid.

Another consideration can be made on the dimension of the task space. Above we argued that the task space could have up to *n*_*ts*_ = 6 degrees of freedom. However, our human body can interact with the external environment at multiple locations i.e., we can use different parts of our body to perform motor tasks. For example, we can use one of our hands to manipulate chopsticks, while using the other hand to hold or move a cup of coffee. In this case, the dimensionality of the task space is larger than 6. A growing dimension of the task-space will shrink the null-space of selectable synergies, thus limiting the hyper-space that is typically available. The limit case is when the task-space dimension reaches the joint-space dimension, thus leading to the absence of a null-space. In other words, the selectable synergies will shrink to only one obligatory synergy. As an example readers may imagine the game of “Twister”, where participants are invited to place their hands and feet in particular configurations, up to a limit where they will find it impossible to complete the task (Foley & Aftorney, 1966).

Given the proposed mechanical framework for describing motor synergies, we can conclude that the number of synergies is determined by the size of the null-space – which can have infinitely many solutions – and not by the number of features (muscles, joints, or task-space variables). This is consistent with the first proposition of the expansion hypothesis: “*the number of synergies is not limited by the number of controlled features*”.

A direct consequence of the first proposition is that – by having infinitely many solutions for the same motor task – different humans might exhibit different synergies to achieve the same goal. The numerical example presented later (Numerical Example Section) shows how this repertoire of different synergies can be generated even for a relatively simple motor task involving a few muscles and joints.

### Developmental Proposition

If we look at the development of motor skills during infancy and childhood, we discover that healthy individuals follow a highly-stereotyped sequence. Seefeldt and Gallahue’s early works provide a good introduction to the different phases of motor development (Gallahue, 1982; Seefeldt, 1980, 1986). At least three main phases can typically be distinguished. The first phase is dominated by reflexes and it characterizes the neonatal and sometimes pre-natal period. Examples are palmar grasp, walking or parachute reflexes to cite some (Anekar & Bordoni, 2022; Cupute et al., 1984; McGraw, 1932). These reflexes involve the coordinated and ‘hard-wired’ motion of multiple muscles and joints. The immediately succeeding phase consists of the inhibition of such reflexes to build what are known as fundamental or rudimentary motor skills. Common examples are crawling, pulling, pushing, throwing, or walking. This phase starts around the first year of age and continues through all of early childhood (1 to 7 years old approximately). In the fundamental motor skill learning phase, children learn to perform voluntary coordinated motor actions necessary to navigate the surrounding environment. The last stage is the so called specific or specialized phase during which children learn more advanced motor skills linked to task-specific actions. Typical examples are sport, dance or motor skills dedicated to musical instruments.

Similar observations but from a different standpoint were drawn by the comprehensive review by Prof. Hadders-Algra. In her work, the Neuronal Group Selection Theory is proposed to explain the development of motor coordination (Hadders-Algra, 2018). One key element of this theory divides the motor development in two phases: the primary variability phase and the secondary variability phase. The former characterizes the pre-natal and neonatal phase in which humans explore their motor functions without adaptation (this could be considered a parallel to the reflexive phase previously presented). The latter, instead, characterizes the effective motor learning phase in which afferent information is used to refine, adapt and build motor skills. This second phase could be matched with the voluntary motor learning phase previously postulated. In the review, it is emphasized how humans start to perform movements even in the pre-natal phase – as early as 7 weeks and 2 days – in correspondence with the development of spinal synapses. In such a phase, fetuses’ movements encompass all body parts; they are slow, small, simple and highly stereotyped. Starting from this stage, the movements start to refine following a gradient from generalization to specialization. Interestingly, such primitive crude coordinated movements have been observed also in other animal species such as chick embryos and fetal guinea pigs (Hamburger, 1973; van Kan et al., 2009). The author uses the term ‘neural coalitions’ to describe groups of neural pathways that allow for increasingly complex ‘elementary actions’. She claims that humans build a repertoire of such neural coalitions as pre-programmed motor solutions for commonly encountered situations. This suggests that the stages of motor development highlight an intriguing aspect: humans consistently progress from crudely coordinated and unspecialized motor actions to finely coordinated and specialized actions, in line with the developmental proposition of the synergy expansion hypothesis.

An example of this crude-coordination-to-specialization behavior can be observed in the literature on the development of prehension. Starting from the pioneering work of Halverson (Halverson, 1931), and followed by the improvements of Hohlstein and Butterworth (Butterworth et al., 1997; Hohlstein, 1982); the literature agrees that prehension follows 3 fundamental steps after the inhibition of the palmar grasp reflex (Anekar & Bordoni, 2022; Niezgoda & Beers, 2017). The first step is characterized by the use of the whole hand in an unspecialized manner. This is followed by a second stage during which children use parts of the hand while some specialization begins to develop. The last phase is reached when infants start using the pads of the distal phalanges, the tips of the fingers, in a specialized manner. In other words, humans proceed from highly coordinated power grasps to finger-specific precision grasps.

This trend, that crude coordination is progressively refined, or developed, in more specialized control is observed not only during childhood, but also in adults when learning more advanced motor skills. An example can be found in piano playing. In the work by Furuya et al., the authors observed how musically naïve subjects acquired individuated finger control after practicing some elementary piano exercises (Furuya et al., 2014). Similar results were observed in a related study by Kimoto et al. in which trained and untrained musicians’ hand dexterity were compared. The results indicated an expertise-dependent increase in finger dexterity and movement independence between the fingers (Kimoto et al., 2019).

From the perspective of the synergy expansion hypothesis, humans might start with a predefined set of crude motor synergies – such as the generalized movements described by Hadders-Algra – e.g., grasping, pulling, pushing, stepping, etc. These **preliminary or primordial synergies** are later refined, fragmented and/or combined to acquire new and more advanced motor skills such as playing a musical instrument, dancing, or practicing a sport. While individuals might start from the same set of primordial synergies, no constraints are placed on how different subjects converge to a given motor action. This is in line with a recent experiment presented by Garcia et al. where they showed how long-term learning in rodents can be systematically represented by multiple transitions through a hierarchy of saddle points (Garcia et al., 2023). The Numerical Example Section provides a possible mathematical description of how multiple subjects, despite starting from the same initial condition, might converge to substantially different synergies.

This does not preclude the use or existence of individual muscle or joint elementary actions i.e., synergy composition vectors (***w***_***i***_) where only one feature is activated – one vector element different from zero – and all the other features are kept inactivated – all the other vector’s elements are zero. Those are possible and could be achieved in healthy unimpaired adult human beings. However, these individual actions appear plausibly as a consequence of the refinement of a previously highly coordinated movement. To state it in a different way, **highly individuated muscle or joint movements can be interpreted as very refined motor synergies** where the coordinated action involves only a small subset (as few as one element) of muscles or joints. Interestingly, evidence of unique activity patterns representing specific movements has been found in human neuroimaging when learning multiple skilled movements, thus supporting the synergy specialization gradient claim (Wiestler & Diedrichsen, 2013).

However, the human musculoskeletal system also presents physical constraints (e.g., tendons and ligaments) that do not allow a perfect uncoupling of every muscle and every joint. This provides a biomechanical limit to the level of individuation that a motor synergy can reach. Todorov and Ghahramani highlighted this in their study on kinematic synergies of the hand showing that synergistic i.e., coordinated, behavior emerged even when subjects where tasked to perform highly individuated joint motions (Todorov & Ghahramani, 2004). Moreover, synergistic behavior might also emerge from the intrinsic constraints of a particular motor task forcing the individual to move certain muscles or joints with a certain configuration e.g., when trying to grasp an object in a cluttered space you adapt arm configuration to avoid obstacles.

An interesting connection may be drawn between the coordination to individuation of motor synergies and the emergence of the unwanted synergies in neurally impaired subjects. For example, post-stroke survivors often show the emergence of undesired coupled movements that won’t allow them to individually move their limbs e.g., the shoulder abduction – elbow flexion synergy (Twitchell, 1951). Could it be that neural damage, and in particular stroke, forces subjects to regress towards the primordial synergies they learned to refine as children, and that motor recovery is a form of re-wiring and re-learning of the more refined synergies of adulthood? Another example might come from the study of posture in stroke-survivors. Boehm et al. showed how stroke survivors tend to change the orientation of their ground reaction force towards a more leaning-forward posture (Boehm & Gruben, 2016), which might be associated to the forward-oriented upright posture of infants.

### Behavioral Proposition

The last aspect of the synergy expansion hypothesis concerns the storing and use of the acquired motor synergies. The authors propose that the learned synergies are stored in memory and recruited upon need. This requires the capability of humans to store specific groups of coordinated actions and re-evoke them. If this is true, we should expect minimal computational effort to perform a learned elementary action, since the human motor controller should ‘simply’ perform a *select-and-shoot strategy* of the necessary synergy required for the task (Figure 4).

**Figure 4.**
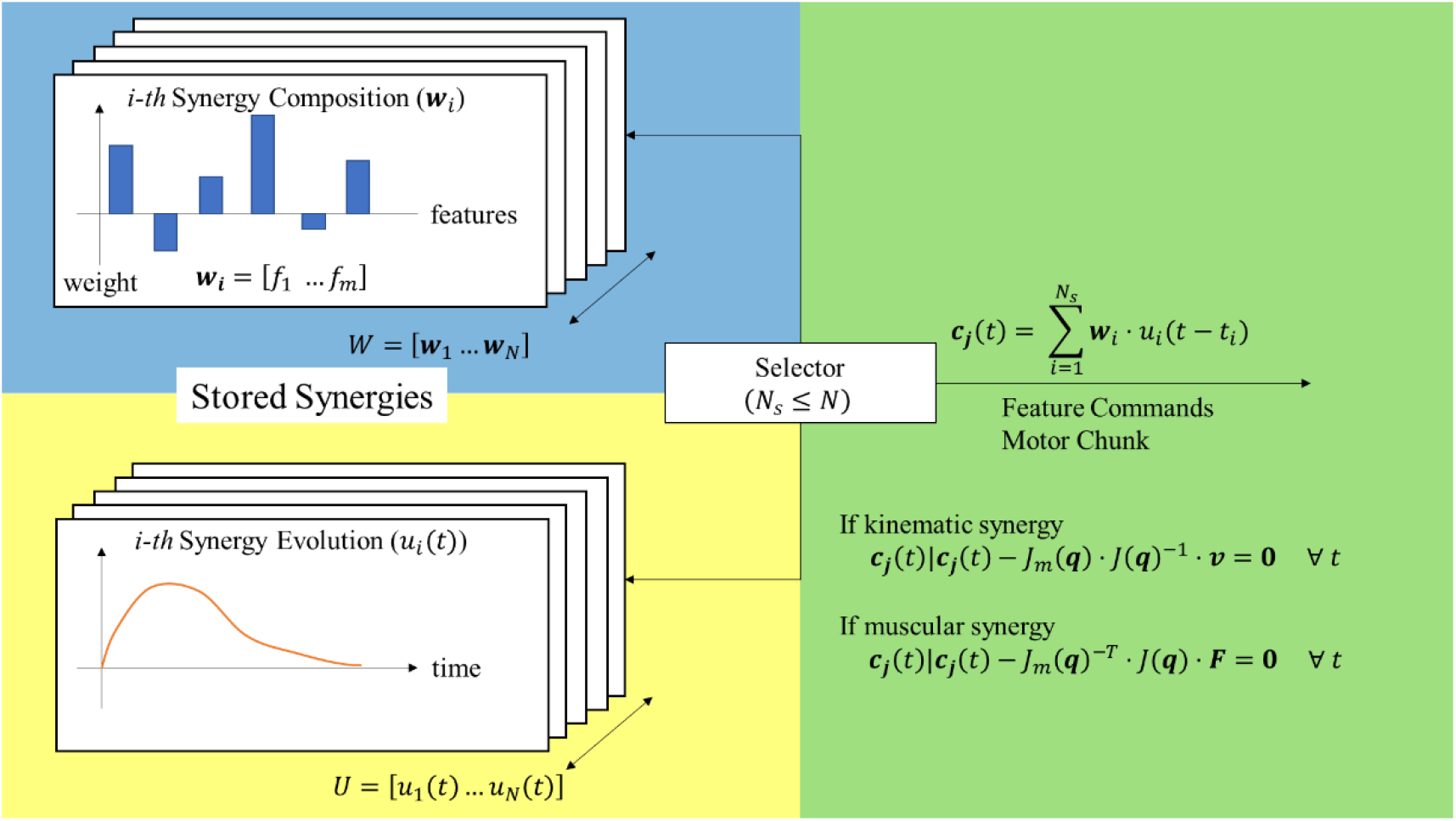
Diagram showing the ‘select-and-shoot’ strategy to generate the feature commands i.e., the motor chunks, to perform a given motor action.

Considering the hierarchical framework presented in Figure 1, we can imagine synergies to be stored as two different elements: (i) the synergy composition vector (***w***_*i*_), a finite-dimensional vector containing the contribution of each controlled feature (*f*) to the considered synergy such that ***w***_***i***_ = [*f*_1_ … *f*_*m*_], where “m” is the number of features controlled by that synergy and “i” is the specific synergy; and (ii) the temporal evolution trajectory (*u*_*i*_(*t*)), a scalar function specifying how a selected synergy composition will evolve over time. As highlighted by the mechanical proposition of the expansion hypothesis, the number of stored synergy compositions *W* = [***w***_**1**_ … ***w***_***N***_] can be greater than the number of controlled features i.e., *N* ≫ *m*. Analogously to the synergy composition, no theoretical limits are imposed to the number of temporal evolutions i.e., *U* = [*u*_1_(*t*) … *u*_*N*_(*t*)] with *N* > 0.

This subdivision of synergies into compositions and temporal evolutions is in line with the recent work proposed by Cheung and Seki, where they proposed a similar architecture for the neural basis of motor synergies (Cheung & Seki, 2021).

When an individual is asked to perform a certain motor task, this will be decomposed into a sequence of motor chunks, which, in turn will be decomposed into a series of elementary actions (refer to Figure 1). At this point, for each elementary action, a “selector” will pick from the stored synergies a certain subset *N*_*s*_ of synergy compositions and temporal evolutions with *N*_*s*_ ≤ *N*. Once selected, the composition and temporal evolutions will be combined to generate the desired feature commands i.e., the desired motor chunk ***c***_***j***_(*t*) – please refer to Equation 1:

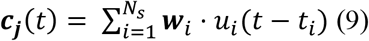

Depending on the type of controlled feature, the resulting commands will have to obey either Equation 7 for kinematic synergies or Equation 8 for muscular synergies.

No particular constraints are placed on the mathematical properties of the synergy composition vector ***w***_***i***_. Nonetheless, an interesting question is whether the vector components are constant or time-varying. This would not affect the ‘select-and-shoot’ strategy, but it would definitely change the role of the synergy evolution component *u*_*i*_(*t* − *t*_*i*_). In other words, if the elements of ***w***_***i***_ can change sufficiently rapidly over time, the temporal evolution terms can be directly included within the synergy composition. The literature on synergy extraction mostly focused on synergy compositions with constant components. However, D’avella and Tresch showed the possibility to decompose muscle patterns also using time-varying synergies (D’avella & Tresch, 2001). This alternative hypothesis is – however – still debated since Hart and Giszter convincingly demonstrated that time-varying synergies cannot account for their data on muscular synergies in frogs and rats (Hart & Giszter, 2013). Another interesting possibility is that the synergy composition vector can be updated over longer time scales to account for motor learning.

In the literature, the synergy evolution patterns *u*_*i*_(*t* − *t*_*i*_) are typically extracted by means of dimensionality reduction, thus reflecting the observed trajectories in the sampled motor tasks. When moving from analysis to synthesis of synergies, no guidelines are available on how to shape such time paths. However, multiple studies on human unconstrained motions highlighted the tendency of humans to move following smooth and continuous trajectories, characterized by bell-shaped velocity profiles (Flash & Hogan, 1985; Hogan, 1985; Morasso, 1981; Mussa-Ivaldi et al., 1985). An interesting study, still not fully explored, would be to investigate the properties of the synergy evolution function *u*_*i*_(*t* − *t*_*i*_), to validate the extension of previous motor control studies to control with motor synergies.

The ‘select-and-shoot’ strategy aligns with the idea of feed-forward predictive motor commands. This is in line with the work by Yeo et al. in which they showed that feed-forward motor commands are necessary to deal with the presence of substantial feedback delay (Yeo et al., 2016). Nonetheless, this does not preclude the presence of feedback control. The literature has widely investigated this possibility, showing how such feedback control models can competently describe many aspects of human motor behavior (Diedrichsen et al., 2010; Kumar et al., 2016; Scott, 2004; Todorov, 2004, 2005; Van Wouwe et al., 2022). In the proposed behavioral proposition, feedback can play a role in adapting – e.g., by means of a sensory driven gain – the synergy composition (***w***_***i***_) or the synergy temporal evolution (*u*_*i*_(*t* − *t*_*i*_)). For example, imagine a case in which a subject is tasked to grasp an object. According to the ‘select-and-shoot’ strategy, the subject will recruit the ‘most adequate’ synergy composition and evolution to perform the task and send out that command (the feed-forward part). However, it is a common tendency of humans to superimpose corrective actions near the end of a reach to a target (Woodworth, 1899). This will require some sort of feedback, and that could be achieved either by turning off the ‘magnitude’ of the previously shot synergy, or by ‘shooting’ a compensating synergy to augment the previous one; the latter seems more plausible (Flash & Henis, 1991).

The selection of the appropriate synergy compositions and evolutions for a given motor task is an interesting question, though outside of the scope of this work. A possible approach can be found in the recent work by Heald et al. in which they propose contextual inference as a way to learn motor repertoires (Heald et al., 2021).

The idea of learning motor skills through a hierarchical process of selection and execution was also recently reviewed by Diedrichsen and Kornysheva (Diedrichsen & Kornysheva, 2015). In their work, the authors provide a theoretical framework to explain motor skill learning through specialized neural circuits. Such circuits are used to produce the required movements in a stable and invariant way. Specifically, the authors re-propose the theory of motor chunking presented by Lashley in 1951 (Lashley, 1951), in which motor chunks represent specific temporal combinations of elementary actions that could be quickly retrieved and accurately combined together. Combination of motor chunks generate a motor sequence that is necessary to complete a certain motor task (refer to Figure 1). The synergies proposed in our work can be considered the building blocks of the elementary actions i.e., specialized neural circuits that are used to ‘select-and-shoot’ the desired movements.

Related work that could help to address the selection process fall within the family of research interested in the processes of ‘motor learning’ and ‘motor adaptation’ widely investigated in multiple neuroscientific works (Brashers-Krug et al., 1996; Heald et al., 2021; Oh & Schweighofer, 2019; Stefan et al., 2005).

### A Numerical Example

An example of how synergies can emerge from the “mechanical proposition” of the expansion hypothesis can be obtained by considering a 3-bar linkage model representing planar motion of the upper-limb as presented in Figure 5.

**Figure 5.**
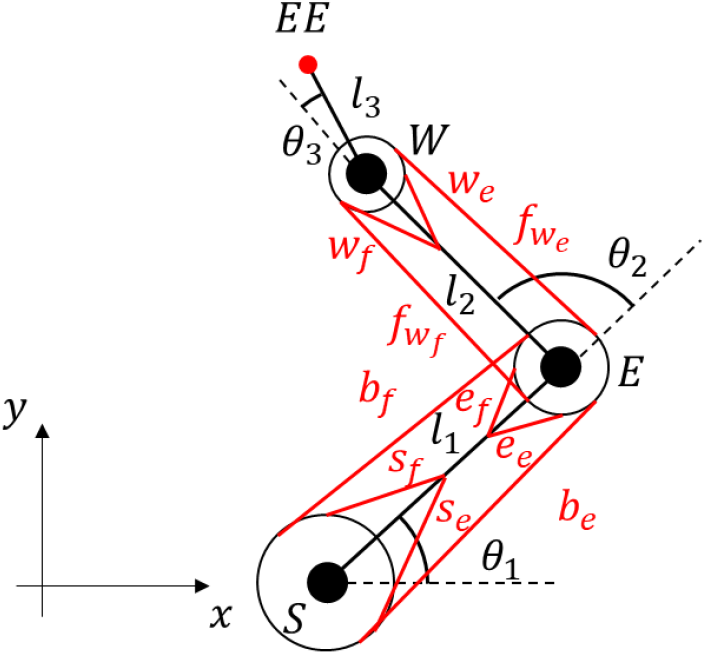
Three bar link model representing planar motion of the upper-limb. ‘S’ represents the shoulder joint, ‘E’ the elbow, ‘W’ the wrist, and ‘EE’ the end-effector. The solid black lines (*l*_1_, *l*_2_, *l*_3_) represent respectively the upper-arm, fore-arm, and hand. The angles (*θ*_1_, *θ*_2_, *θ*_3_) are respectively the shoulder, elbow, and wrist joint angles. The solid red lines represent the line of action of 10 muscles: two shoulder mono-articular flexors (*s*_*f*_, *s*_*e*_), two elbow mono-articular flexors (*e*_*f*_, *e*_*e*_), two wrist mono-articular flexors (*w*_*f*_, *w*_*e*_), two bi-articular shoulder-elbow flexors (*b*_*f*_, *b*_*e*_), two mono-articular fore-arm wrist flexors 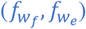.

Considering the joint relative angle coordinates (*θ*_1_, *θ*_2_, *θ*_3_) as configuration variables, the end-effector (EE) task-space kinematics can be described as:

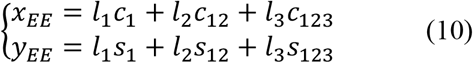

where *c*_*i*_, *s*_*i*_ are notations respectively for the cosine and sine functions e.g., *c*_12_ = cos(*θ*_1_ + *θ*_2_); while *l*_*i*_ are the lengths of the rigid links representing respectively the upper-arm, fore-arm, and hand.

The task-space Jacobian is obtained from the task-space kinematics through partial differentiation of Eqs. (10):

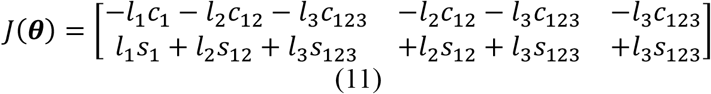

Similarly, the joint relative angle coordinates can be used as configuration variables to describe muscle shortening. In the case considered, we focused on a 10-muscle system composed of: 2 mono-articular shoulder flexor-extensors of length (*s*_*f*_, *s*_*e*_), 2 mono-articular elbow flexor-extensors of length (*e*_*f*_, *e*_*e*_), 2 mono-articular wrist flexor-extensors of length (*w*_*f*_, *w*_*e*_), 2 bi-articular shoulder-elbow flexor-extensors of length (*b*_*f*_, *b*_*e*_), and 2 mono-articular forearm wrist flexor-extensor 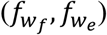. The relation mapping joint angles to muscle contractions is uniquely defined by geometry:

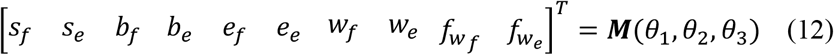

where ***M*** is a vectorial function ***M*** = [*M*_1_ … *M*_10_], with *M*_*i*_: ℝ^3^ → ℝ^10^, that maps the joint angular trajectories to the muscle elongations. By means of partial differentiation of ***M*** with respect to *θ*_*i*_, the muscle Jacobian can be computed as:

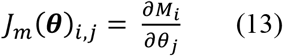

Two Jacobian (*J, J*_*m*_) matrices can be used to move, respectively, from the two-dimensional task space (*x, y*) to the three-dimensional joint space (*θ*_1_, *θ*_2_, *θ*_3_), and from that joint space to the ten-dimensional muscle space 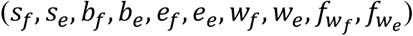.

Consider now the case in which we are tasked to generate a series of planar forces (*F*_*x*_, *F*_*y*_) at the end-effector – i.e., the hand – of our arm model in a given arm configuration (*θ*_1_, *θ*_2_, *θ*_3_). In order to generate such forces, the shoulder, elbow and wrist joints will have to generate – according to Equation (5) – a set of uniquely defined torques, namely an **obligatory synergy**:

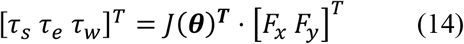

Figure 6 provides a graphical example of the generation of such obligatory synergies by means of Equation 14. Specifically, you can observe that for every end-effector force vector [*F*_*x*_ *F*_*y*_]^*T*^, a uniquely defined torque vector [*τ*_*s*_ *τ*_*e*_ *τ*_*w*_]^*T*^ is obtained.

**Figure 6.**
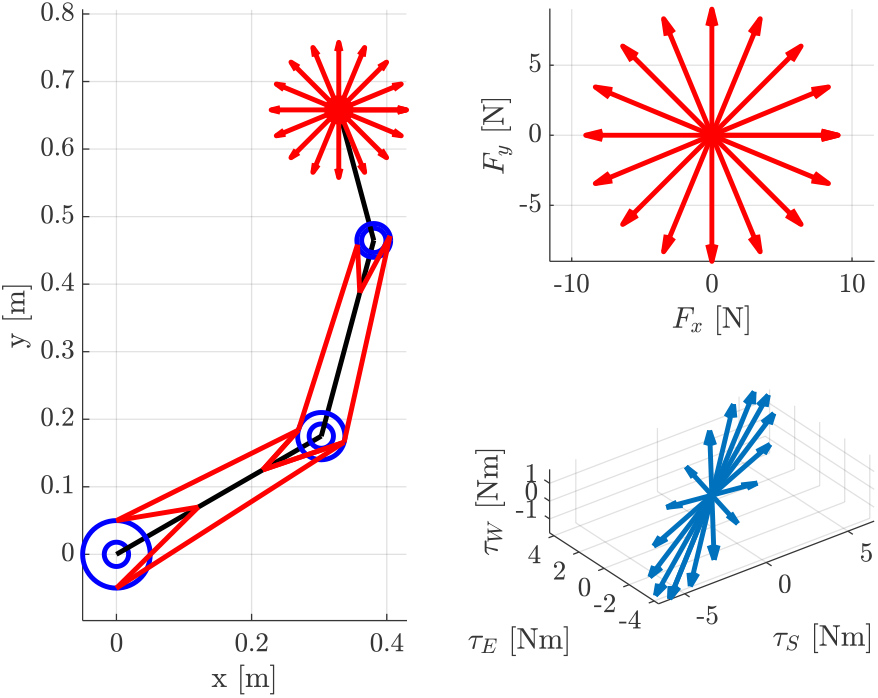
Example of the generation of obligatory synergies in joint-space. The left panel shows the arm model in Cartesian space with a series of end-effector forces represented with red arrows. The right top panel shows the amplitude and direction of such forces in task-space. The right bottom panel presents the mapping of such forces in torque-space.

When moving from joint-space to muscle-space, in order to generate the desired torque coordination (*τ*_*s*_, *τ*_*e*_, *τ*_*w*_), the subject will have to generate a combination of muscle forces in agreement with the inverse of Equation (6):

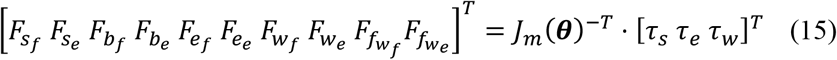

However, as explained in the Mechanical Proposition section, the inversion of *J*_*m*_(***θ***)^*T*^ is not uniquely defined i.e., the system of equations is under-determined. This leaves space for infinitely many combinations of muscle forces to generate the same torque coordination. These are what we defined as **selectable** synergies. In this specific case, the infinitely many combinations will lie in the null-space of *J*_*m*_(***θ***)^*T*^ which will have a dimensionality equal to *n*_*ns*_ ≤ *n*_*m*_ − *n*_*τ*_, where *n*_*ns*_ is the size of the null-space, *n*_*m*_ is the number of muscles and *n*_*τ*_ the number of torques.

A numerical simulation was implemented to show how the same task could be completed using different i.e., **selectable**, synergies. Assume a group of subjects is tasked to learn how to generate a certain arm end-effector force ***F*** in a planar configuration (refer to Figure 7). The previously presented arm model is adopted. For example, consider the generation of 10 N of force in the positive ‘x’ direction.

**Figure 7.**
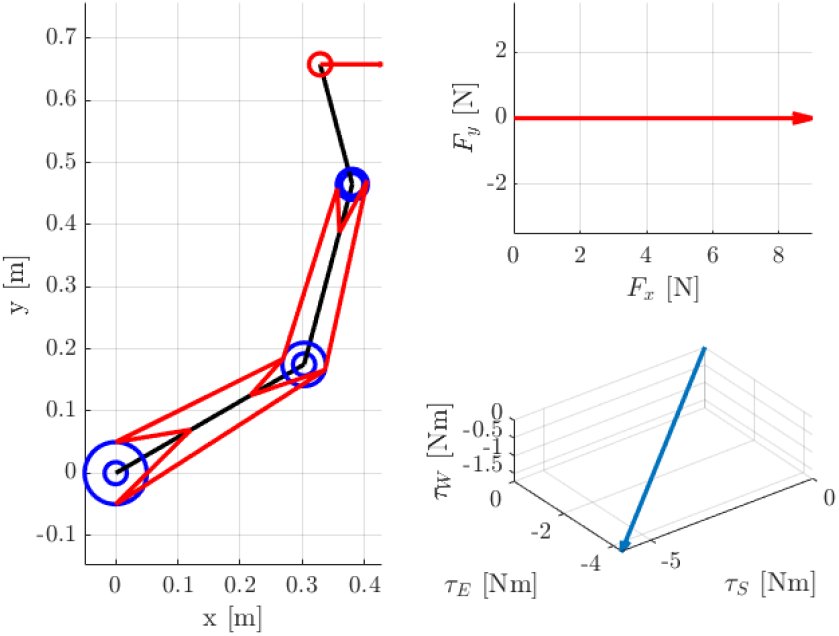
Three body rigid arm model generating a horizontal arm force. The left panel shows the arm configuration and desired force direction (red arrow). The top right panel show the desired end-effector force, while the bottom right panel show the obligatory torque coordination to generate such end-effector force.

Assume also that each subject starts from a rest condition for which ***F***_***m*0**_ = **0** ∈ ℝ^10^. In such a scenario, we can imagine subjects to learn this task by iteratively varying their muscle force vector ***F***_***m***_ to satisfy the following equality:

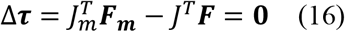

Algorithmically, this may be modeled as a stochastic gradient descent learning problem^1^, where subjects are trying to minimize an error that can be defined – for example – as:

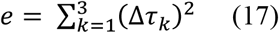

We do not claim that this is exactly how humans learn a new behavior. However, it is a plausible theory. Importantly, by imposing a minimization of task error (Eq. 17), we do not uniquely define joint space configurations, thus leaving space for the emergence of multiple ‘good-enough’ synergies.

At each new iteration ‘i’, a series of 5 new possible force vectors 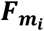 were generated by adding a random increment Δ*F* to each element of 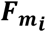. For each of these 5 new possible force vectors, the algorithm selected the one minimizing the error function (Eq. 17). This was done to simulate the exploration of different directions in the muscle force hyper-space. The force vector at the next iteration was equal to:

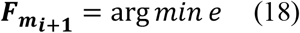

The learning continued until the error went below an arbitrary threshold i.e., *e* ≤ *e*_*th*_. In the simulation, our increment Δ*F* was a pseudo-random uniformly distributed value of range ±25 *N*. However, since muscles can only pull (*F* > 0), if the resulting muscle force was less than zero, the algorithm forced it to zero. A threshold of *e*_*th*_ = 1 · 10^−2^*N*^2^*m*^2^ was chosen. The error function *e* used the sum of squares in order to avoid cancellation due to sign changes.

Alternatively, the absolute value function could have been used.

Figure 8 presents the error function trend for 50 different runs of the proposed learning algorithm, all starting from the same initial zero muscle force vector ***F***_***m***_. All of the runs converged below the imposed threshold – notably at different iterations. This confirms that each of the 50 generated muscle force synergies ***F***_***m***_ was capable of producing the required output force ***F***.

**Figure 8.**
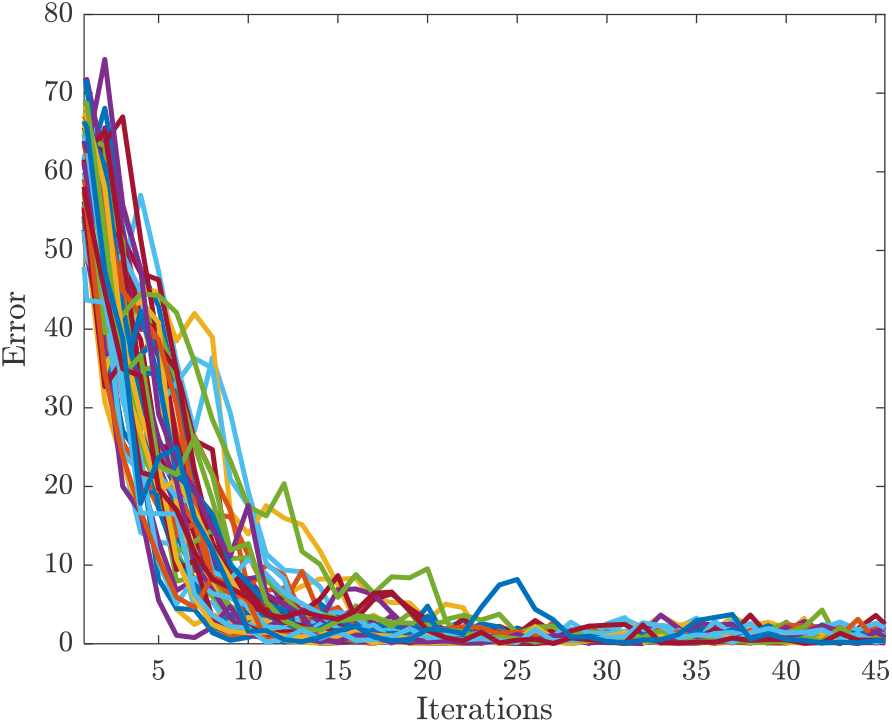
Trend of the error function with respect to the number of iterations for the stochastic gradient descent learning algorithm of selectable muscle force synergies. Each colored line represents a different run of the algorithm.

In addition to the 50 generated selectable muscle synergies, the least-square solution was added to provide an additional comparison. This was computed by solving Equation 15 with a constrained Moore-Penrose inverse (Moore, 1920; Penrose, 1955). The constrained approach was chosen to enforce only positive forces. Despite the fact that all runs started from the same initial condition i.e., resting muscle force, the algorithm converged to diverse solutions. This supports the developmental claim that, although individuals may start with the same set of primordial synergies, there are no constraints on how different subjects may develop different synergies to achieve the same motor action.

An interesting question is how similar or dissimilar these generated muscle synergies were between each other. This was quantified using the Dimension-Insensitive Euclidean Metric (DIEM) proposed by Tessari and Hogan (Tessari & Hogan, 2024). This metric provides a robust measure of the distance between multi-dimensional quantities such as muscle synergies. A DIEM value of 0 means that the distance between two quantities is statistically indistinguishable from that obtained by comparison of randomly generated vectors. Negative values of DIEM indicate similarity, while positive values indicate dissimilarity.

Figure 9 presents the DIEM distribution computed between all the possible combinations of the 51 generated muscle synergies 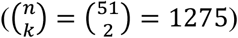. The resulting distribution had a modal value of *DIEM*_*mode*_ = −12.50, and presented a diverse range of similarity/dissimilarity between the 51 generated muscle synergies (−20 < DIEM < +6), thus supporting the claim that there exist multiple distinct solutions to the same motor task.

**Figure 9.**
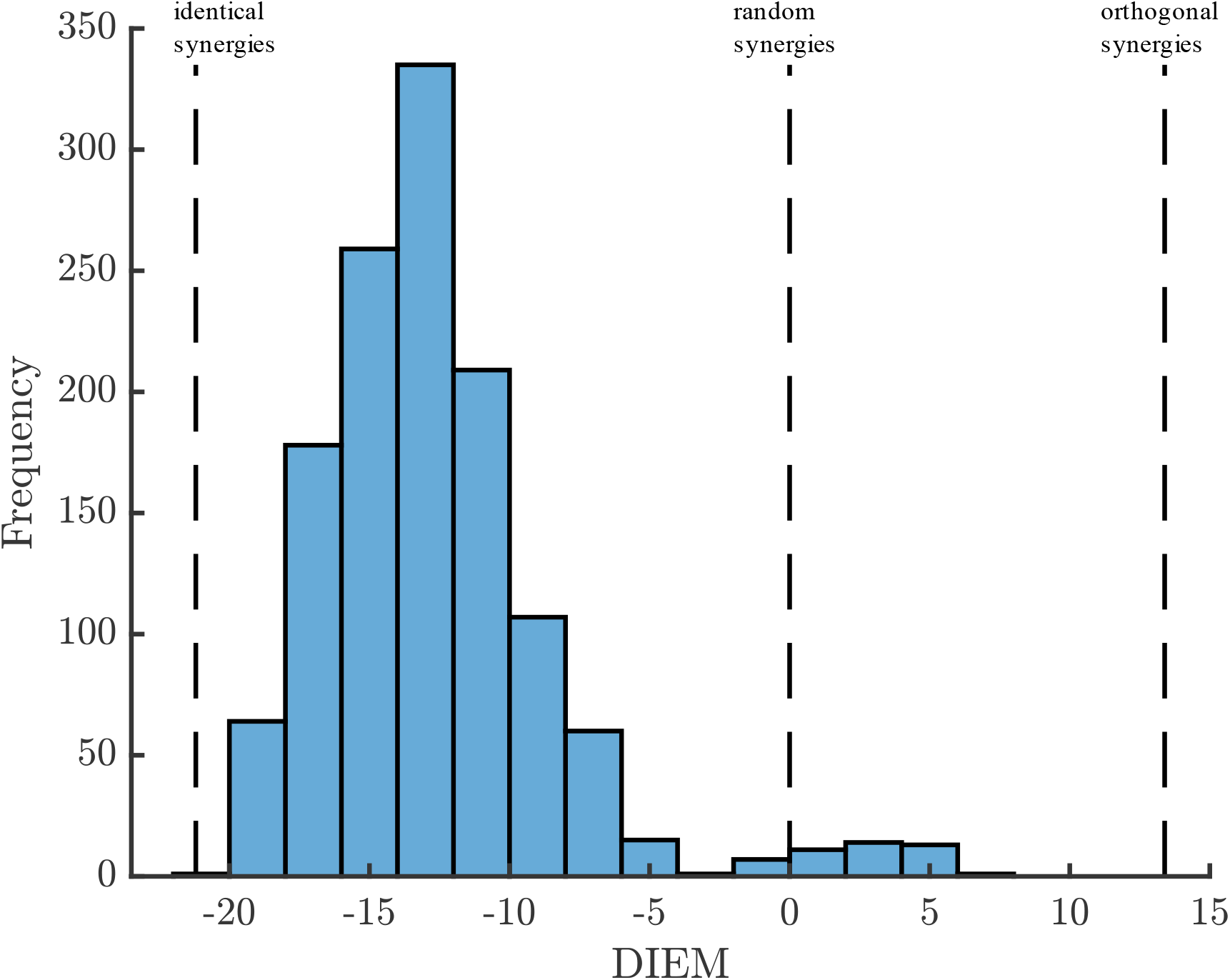
Distribution of the Dimension-Insensitive Euclidean Metric (DIEM) computed between all the 51 different generated muscle synergies. The vertical dashed lines represent the DIEM values respectively for: identical, random, and orthogonal synergies.

Among the 51 generated muscles synergies, Figure 10 reports the most similar synergies, two synergies near the modal value of the DIEM distribution, and the most dissimilar synergies. From Figure 10 we can appreciate that equally successful synergies can assume very different values. In other words, for a bi-dimensional force generation task (planar force) with a 10-dimensional muscle space, it was possible to synthesize 51 different muscle synergies – far greater than the number of muscles (10) – all equally effective to generate the desired task space force.

**Figure 10.**
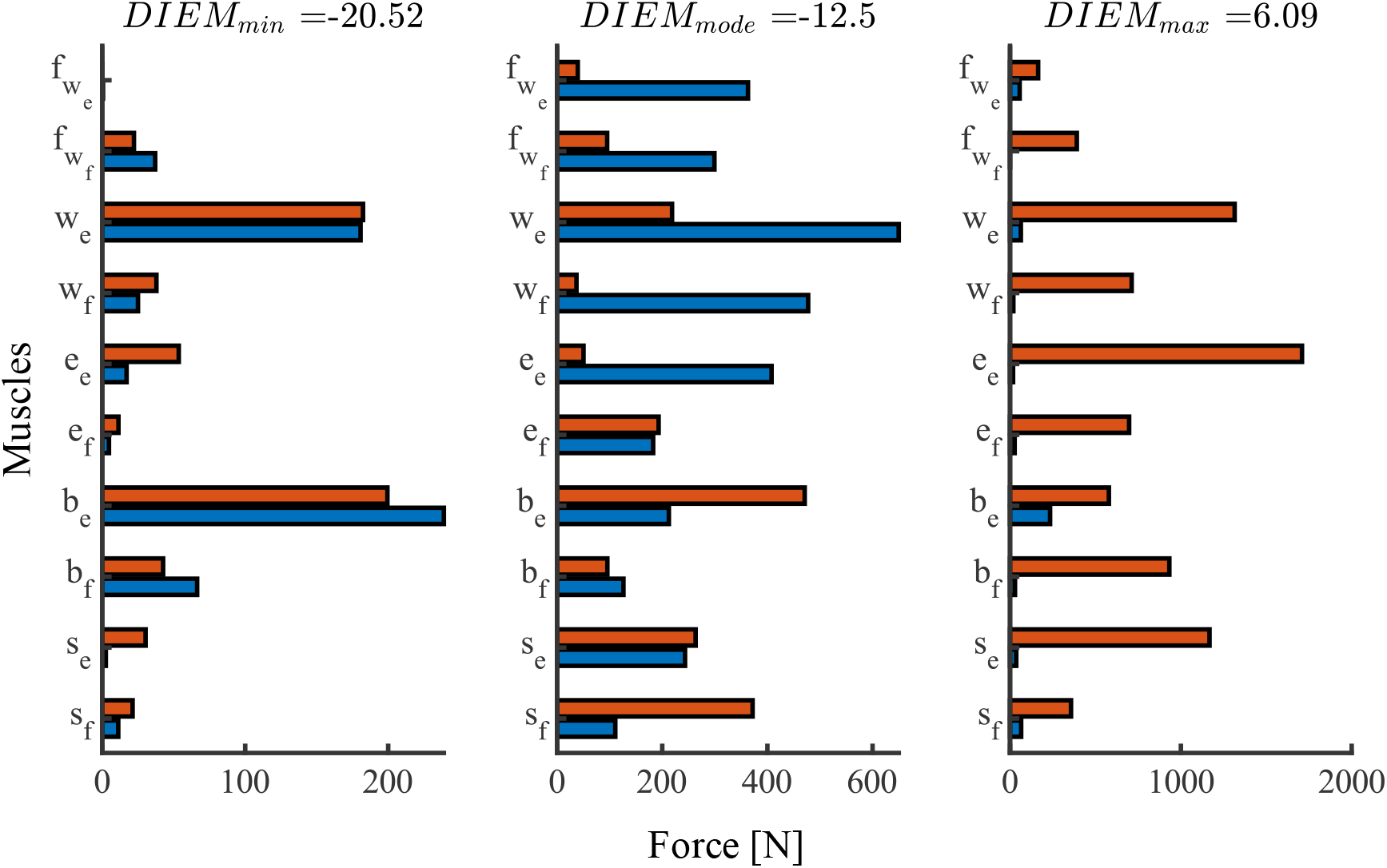
Bar plots presenting the muscles forces of: (left) the two most similar (*DIEM*_*min*_ = −20.52) generated synergies, (center) two synergies near the modal value of the DIEM distribution (*DIEM*_*mode*_ = −12.50), and (right) the two most dissimilar synergies (*DIEM*_*max*_ = 6.09). The y-axis labels represent the 10 different muscles: two shoulder mono-articular flexors (*s*_*f*_, *s*_*e*_), two elbow mono-articular flexors (*e*_*f*_, *e*_*e*_), two wrist mono-articular flexors (*w*_*f*_, *w*_*e*_), two bi-articular shoulder-elbow flexors (*b*_*f*_, *b*_*e*_), two mono-articular fore-arm wrist flexors 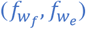.

If we imagine every separate simulation as a different subject during his/her learning of the force generation task, this result confirms that it is possible to reach different solutions to the same motor task, despite starting from the exact same initial condition i.e., primordial motor synergy. In other words, there can be more synergies than the number of features (in our example, muscles). This is consistent with both the Mechanical and Developmental propositions of the expansion hypothesis.

The same numerical analysis has been performed also for three other end-effector force directions: negative ‘x’, positive ‘y’, and negative ‘y’. Similar results were found for all the analyzed cases. The Supplementary Information contains the complete results of these simulations.

This numerical example – focused on the generation of end-effector forces at a constant planar arm configuration – showed the emergence of multiple **selectable** muscle synergies to deliver the same end-effector force. However, this approach can be generalized to kinematic synergies, as explained in the Mechanical Proposition section.

## Discussion & Conclusion

In this work we proposed a new framework for the synthesis and use of motor synergies to answer Bernstein’s coordination problem. The key idea is that motor synergies are the building blocks of human motor control that emerge as a natural consequence of the motor ‘abundance’ of our musculoskeletal system (Mechanical Proposition). These synergies are the result of motor development as it appears that humans learn motor skills starting from crudely coordinated patterns – primordial synergies in which almost all features exhibit the same time course (e.g., the fingers in a power grasp)– and continuously expand their motor skills to highly specialized actions (Developmental Proposition). Learned synergies are stored/memorized and evoked upon need following theories of ‘motor learning’ and ‘adaptation’ (Behavioral Proposition).

As of today, motor synergies have been considered to be an approach to simplifying control that humans might employ to manage their highly complex musculoskeletal system. This is consistent with the proposed definition of a synergy – *the coordinated and simultaneous activity of a finite group of features (neural, muscular, kinematic) to perform a specific motor task*. The expansion framework fully agrees with this vision. The fact that we can have infinitely many synergies, many more than the number of controlled features, simply highlights that there is no upper limit to the number of synergies that may exist. However, it does not contradict the dimensionality reduction purpose of some synergies.

When moving from primordial synergies to highly specialized ones, no claims were here made about whether primordial synergies are substituted by specialized ones or new synergies are generated and added to the library of available commands. However, the work by Yokoi and Diedrichsen suggests that both coordinated actions (sequences of movements) and individuated actions (individual finger movements) are stored in different locations of the central nervous system (Yokoi & Diedrichsen, 2019). This evidence suggests that specialized synergies did not replace crudely coordinated synergies, but were added to the existing repertoire of possible elementary actions. This claim also finds support in the pioneering work by Dominici et al. in which they investigated the evolution of stepping in different stages of human development as well as in other animal models (Dominici et al., 2011). Surprisingly, their study showed common muscular synergies between human babies, toddlers and adults as well other animals. Such common synergies can be associated to the ‘primordial synergies’ presented in the second proposition of the expansion hypothesis: “humans start from highly stereotyped and crude coordinated patterns (primordial synergies) and, through motor learning and development, expand to new and more specialized synergies”. Moreover, the additional stepping muscular synergies observed in adult humans did not substitute for the ‘primordial’ synergy patterns, thus reinforcing the idea that specialization is adding to crude coordination rather than replacing it. Stated differently, this indicates that ‘new’ synergies do not replace ‘old’ synergies but augment them.

It is worth emphasizing that this trend from crude coordination to fine specialization observed in humans may seem counter-intuitive, since the theoretical apex of motor coordination and specialization would culminate in individual joint or muscle movements i.e., a complete absence of synergistic activity. Moreover, it is also important to stress that this behavior in human motor development is the opposite of what robotic engineers typically do. It is common to start programming robotic movements beginning with individual joint motions and later adding coordination and complexity. Is it because the hardware used is simpler and often lacks a null space? Answering the question of why biological systems adopt this reverse approach could be potentially an interesting avenue for future research.

The expansion hypothesis focused on the mechanical spaces (muscle, joint and task) of humans where a mapping between the different domains can be drawn based on geometry and configuration. However, when we move from muscle space to neural space – whether it is spinal or cortical – there is not (to the best of the authors’ knowledge) a clear way to build a continuous mapping between the impressive dimensionality of neural space and muscle space. While the challenge to find such a mapping remains open, several works demonstrated the possibility to decode multiple motor actions from the central nervous system of humans, rodents and primates (Inoue et al., 2018; Kennedy & Schwartz, 2019a, 2019b; Keshavarzi et al., 2022; Moran & Schwartz, 1999; Rodman & Albright, 1987; Schwartz, 1993). Nonetheless, the large disparity between the size of the neural space and the muscle space suggests the presence of a dimensionally-large redundant space within which individuals can find multiple pathways to the same solution. This is further stressed in the recent review by Cheung and Seki in which they tried to tackle the neural basis of motor synergies. Their work highlighted the important role of several factors such as the motor cortex, brain stem and spinal cord in the coordination, temporal activation and tuning of muscle synergies (Cheung & Seki, 2021). They also present a hypothetical hierarchical structure to organize the activation and driving of motor synergies with two distinct neural mechanisms. Their idea – in agreement with previous work on muscle and kinematic synergies (Cheung et al., 2009; D’Avella & Bizzi, 2005; D’avella & Tresch, 2001; ason et al., 2001; Santello et al., 1998, 2016) – is that motor synergies are characterized by two different mechanisms: a synergy organization component that accounts for the contribution of each feature to the considered synergy, and a temporal activation component that informs the temporal evolution of the synergy itself (Giszter & Hart, 2013; Hart & Giszter, 2013).

Such a hierarchical structure is also accepted as a reliable framework to describe the generation of motion. In a hierarchical framework, neural commands – representing sequences of motor chunks, in turn composed by combination of elementary actions – are selected at the cortical level within the premotor area, basal ganglia (Doya, 1999) and striatal centers actively involved in the selection process (Jin et al., 2014).

As explained at the beginning of the paper, elementary actions are proposed as the ‘atomic’ elements of coordinated motion. Each elementary action is defined by a unique temporal evolution and a finite-dimensional vector containing the contribution of each feature (muscle or joint) to that elementary action. A chain of elementary actions generates a motor chunk, which is a temporally bounded activity.

Once selected, these actions are then executed using circuits such as central pattern generators, which are typically located in the spinal cord and the brain stem (Brown, 1914; Churchland et al., 2012; Dominici et al., 2011; Marder & Bucher, 2001). This neural organization of motor commands aligns well with what we propose in the ‘behavioral’ proposition of the expansion hypothesis i.e., the ‘select-and-shoot’ strategy.

The extreme redundancy of the human motor control system was also approached by the theory of dimensional ‘bliss’ and motor abundance introduced by Latash (Latash, 2012). He suggested that human redundancy allows the central nervous system to generate multiple solutions to the same task – aligning with Bernstein’s classical observation of “repetition without repetition” (Bernstein, 1967). The expansion hypothesis agrees with these observations, suggesting that – at least at the muscular and joint level – the availability of these multiple solutions is ensured by mechanics.

The expansion hypothesis focused on the process of evolution, number and selection of features (synergy composition vector ***w***_*i*_) within a given elementary action. However, no specific claims were made on how the temporal evolutions, and in general the temporal execution of the task (chunking and sequence) should happen. This goes beyond the scope of this work, and it might be an interesting future avenue of exploration.

Experiments should be developed to test and validate the expansion hypothesis, with a particular emphasis on: (i) the availability of more synergies than the number of controlled features (mechanical), (ii) the refinement of synergies from coordination to individuation (developmental), and (iii) the possibility to recruit and activate synergies to perform motor tasks (behavioral). The authors recommend especially two experiments. A first one, to test the ‘ echanical’ Proposition, should look at the differences between subjects in performing the same set of motor actions. How similar or different are the observed synergies? Is it possible to find more synergies than the number of controlled features?

A second experiment, to test the ‘Developmental’ Proposition, should look at the number of synergies (either muscular or kinematic) during childhood development or during motor learning of specific tasks in adults. Can we observe a growing number of synergies with age or with increased skill? Do the synergies share common features with the primordial ones?

Another interesting aspect worth pursuing in the future is the eventual difference between synergies used for more kinematic oriented motor actions (such as reaching), with synergies used to generate force or stabilize postures e.g., handling a tool. A very recent work by Borzelli et al. has brought up the possibility that different synaptic circuits might be involved in force generation versus stiffness stabilization. Exploring this aspect, might help us better understand the role of different synergies in motor selection and execution (Borzelli et al., 2024).

In conclusion, this paper motivates these interesting topics of future research. The synergy expansion hypothesis is a foundational framework for future research in the field of human neuromotor control with the aim to provide new answers to Bernstein’s – still open – problem.

## Data Availability

All data generated or analyzed during this study are included in this published article and its supplementary information files.

## Acknowledgments

This work has been funded by the Eric P. and Evelyn E. Newman Fund.

## Supplementary Information

### Selectable Synergies in Different Directions

The numerical analysis of the generation of different selectable muscle synergies presented in the main manuscript for a given end-effector force direction (positive ‘x’) was replicated for three additional end-effector force directions: negative ‘x’, positive ‘y’, negative ‘y’. The same force magnitude of 10 was maintained across conditions. 50+1 muscle synergies were generated for each case following the method presented in the Numerical Example Section.

#### Negative ‘x’ Direction

**Figure 11.**
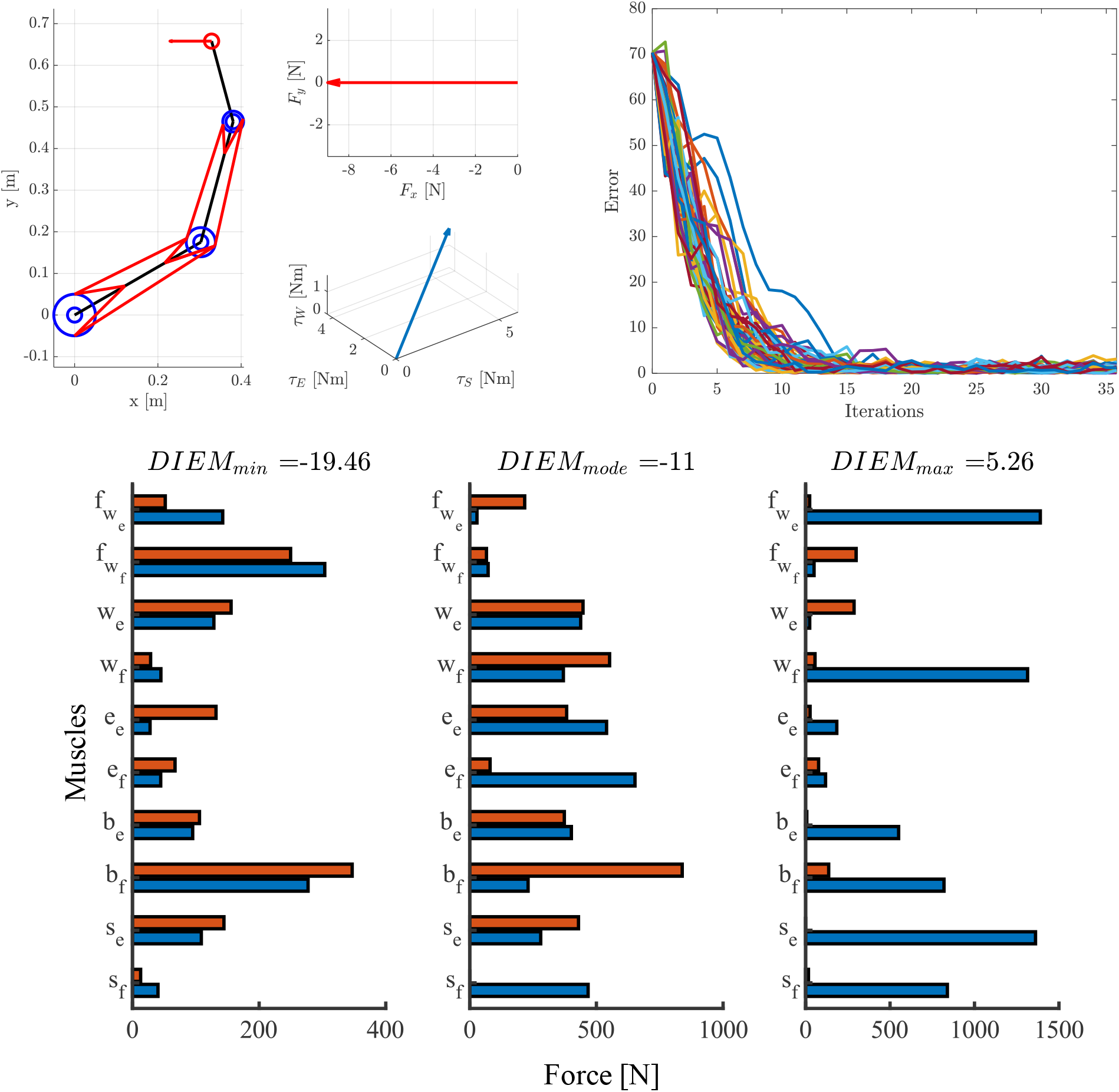
**Top Left**: Three-body rigid arm model generating a horizontal arm force. The left panel shows the arm configuration and desired force direction (red arrow). The top right panel shows the desired end-effector force, while the bottom right panel show the obligatory torque coordination to generate such end -effector force. **Top Right**: Trend of the error function with respect to the number of iterations for the stochastic gradient descent l earning algorithm of selectable muscle force synergies. Each colored line represents a different run of the algorithm. **Bottom**: Bar plots presenting the muscles forces of: (left) the two most similar (*DIEM*_*min*_ = −19.46) generated synergies, (center) two synergies near the modal value of the DIEM distribution (*DIEM*_*mean*_ = −11.00), and (right) the two most dissimilar synergies (*DIEM*_*max*_ = 5.26). These synergies were selected among the 51 different generated synergies. The y -axis labels represent the 10 different muscles: two shoulder mono-articular flexors (*s*_*f*_, *s*_*e*_), two elbow mono-articular flexors (*e*_*f*_, *e*_*e*_), two wrist mono-articular flexors (*w*_*f*_, *w*_*e*_), two bi-articular shoulder-elbow flexors (*b*_*f*_, *b*_*e*_), two mono-articular fore-arm wrist flexors 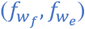.

**Figure 12.**
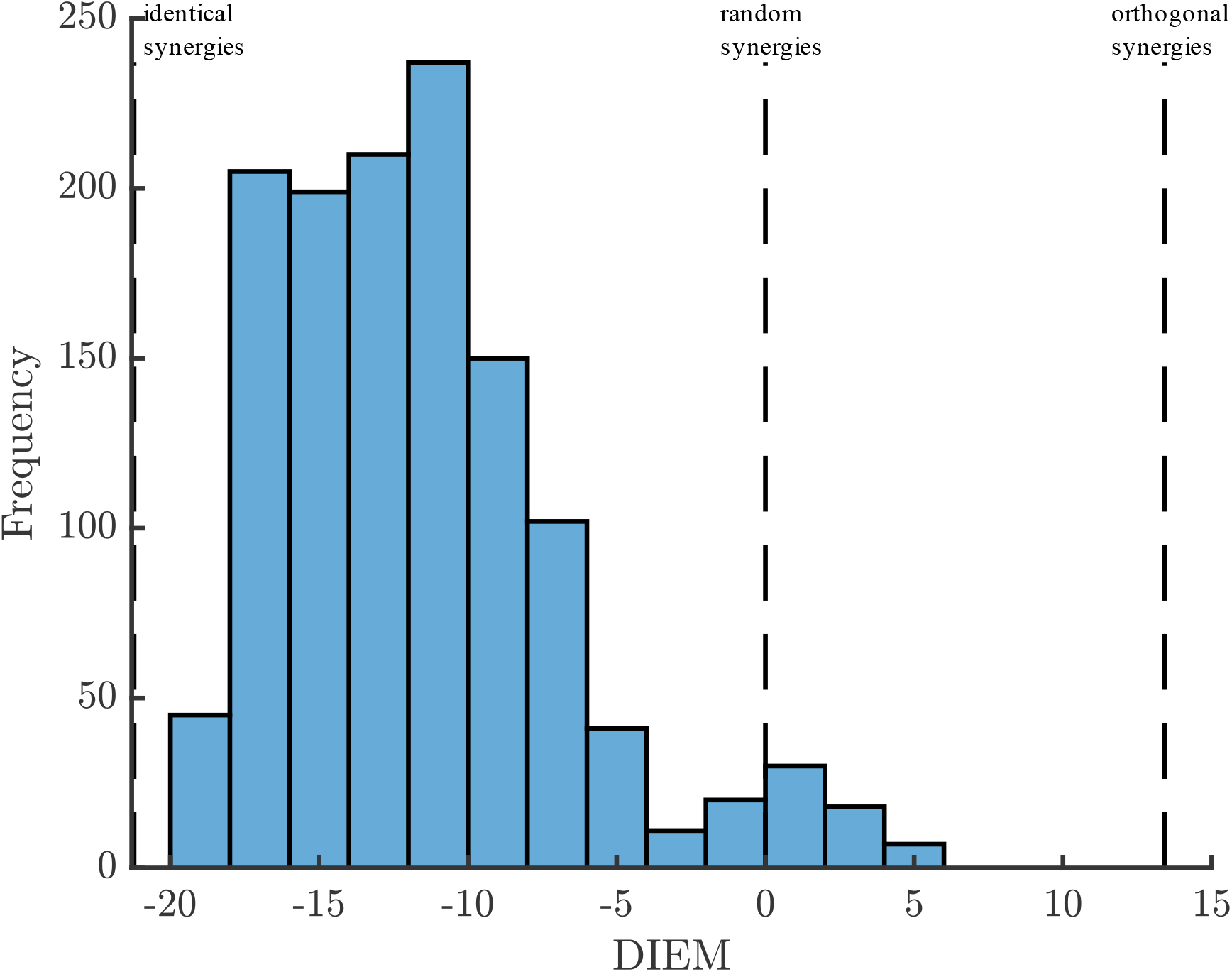
Distribution of the Dimension-Insensitive Euclidean Metric (DIEM) computed between all the 51 different generated muscle synergies. The vertical dashed lines represent the DIEM values respectively for: identical, random, and orthogonal synergies.

#### Positive ‘y’ Direction

**Figure 13.**
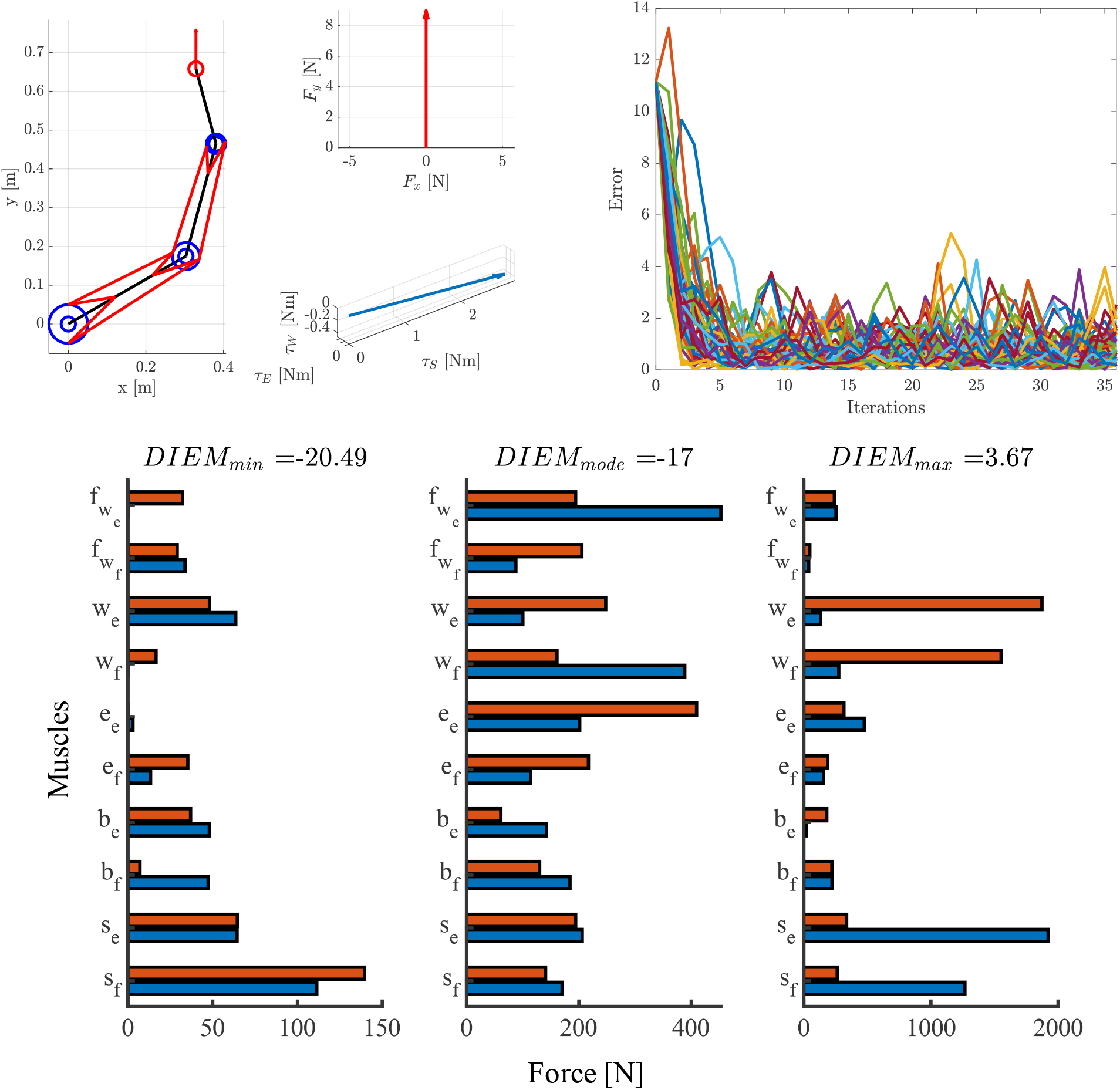
**Top Left**: Three-body rigid arm model generating a horizontal arm force. The left panel shows the arm configuration and desired force direction (red arrow). The top right panel show the desired end -effector force, while the bottom right panel show the obligatory torque coordination to generate such end-effector force. **Top Right**: Trend of the error function with respect to the number of iterations for the stochastic gradient descent learning algorithm of selectable muscle force synergies. Each colored line represents a different run of the algorithm. **Bottom**: Bar plots presenting the muscles forces of: (left) the two most similar (*DIEM*_*min*_ = −20.49) generated synergies, (center) two synergies near the modal value of the DIEM distribution (*DIEM*_*mean*_ = −17.00) synergies, and (right) the two most dissimilar synergies (*DIEM*_*max*_ = 3.67). These synergies were selected among the 51 different generated synergies. The y-axis labels represent the 10 different muscles: two shoulder mono-articular flexors (*s*_*f*_, *s*_*e*_), two elbow mono-articular flexors (*e*_*f*_, *e*_*e*_), two wrist mono-articular flexors (*w*_*f*_, *w*_*e*_), two bi-articular shoulder-elbow flexors (*b*_*f*_, *b*_*e*_), two mono-articular fore-arm wrist flexors 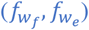.

**Figure 14.**
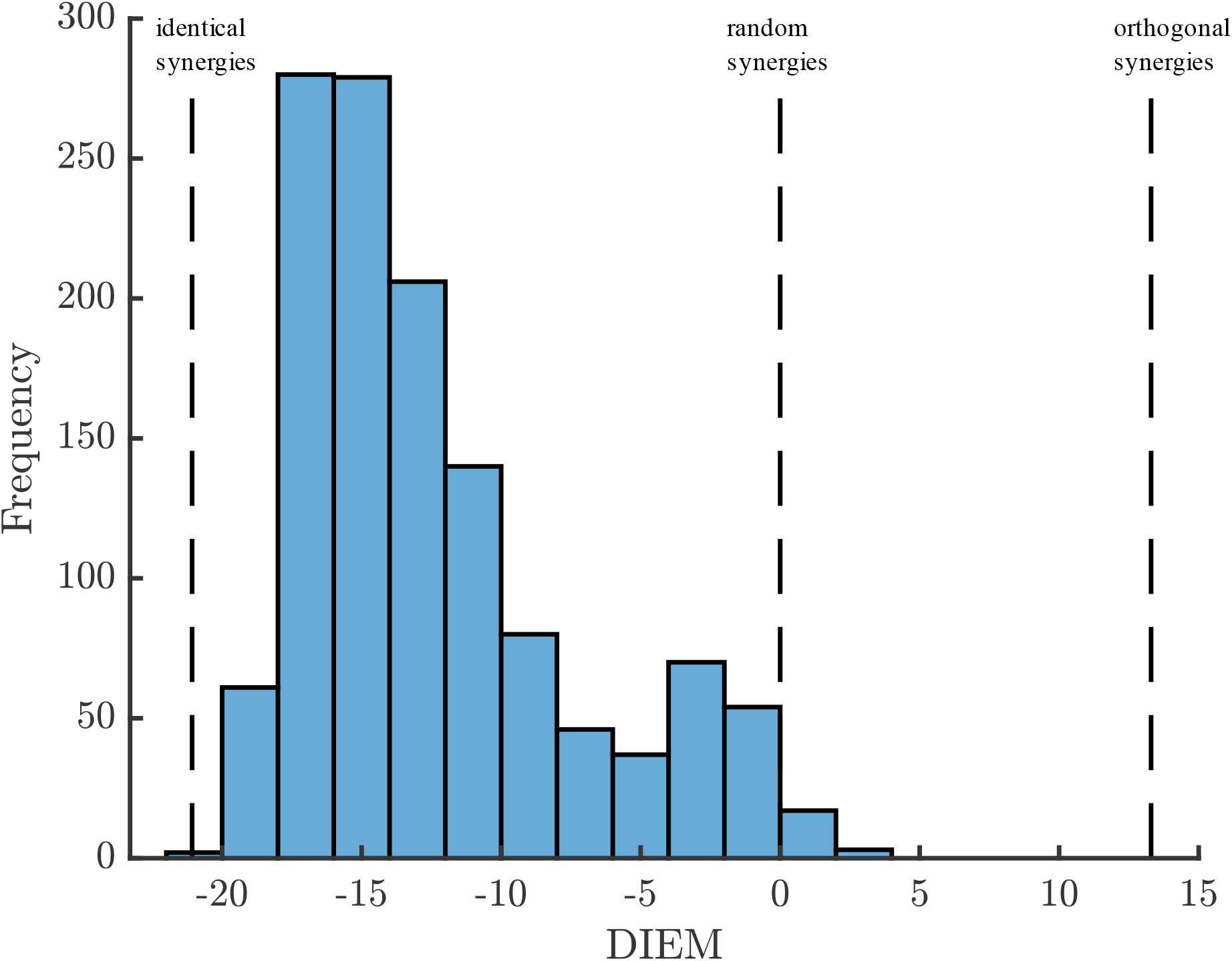
Distribution of the Dimension-Insensitive Euclidean Metric (DIEM) computed between all the 51 different generated muscle synergies. The vertical dashed lines represent the DIEM values respectively for: identical, random, and orthogonal synergies.

#### Negative ‘y’ Direction

**Figure 15.**
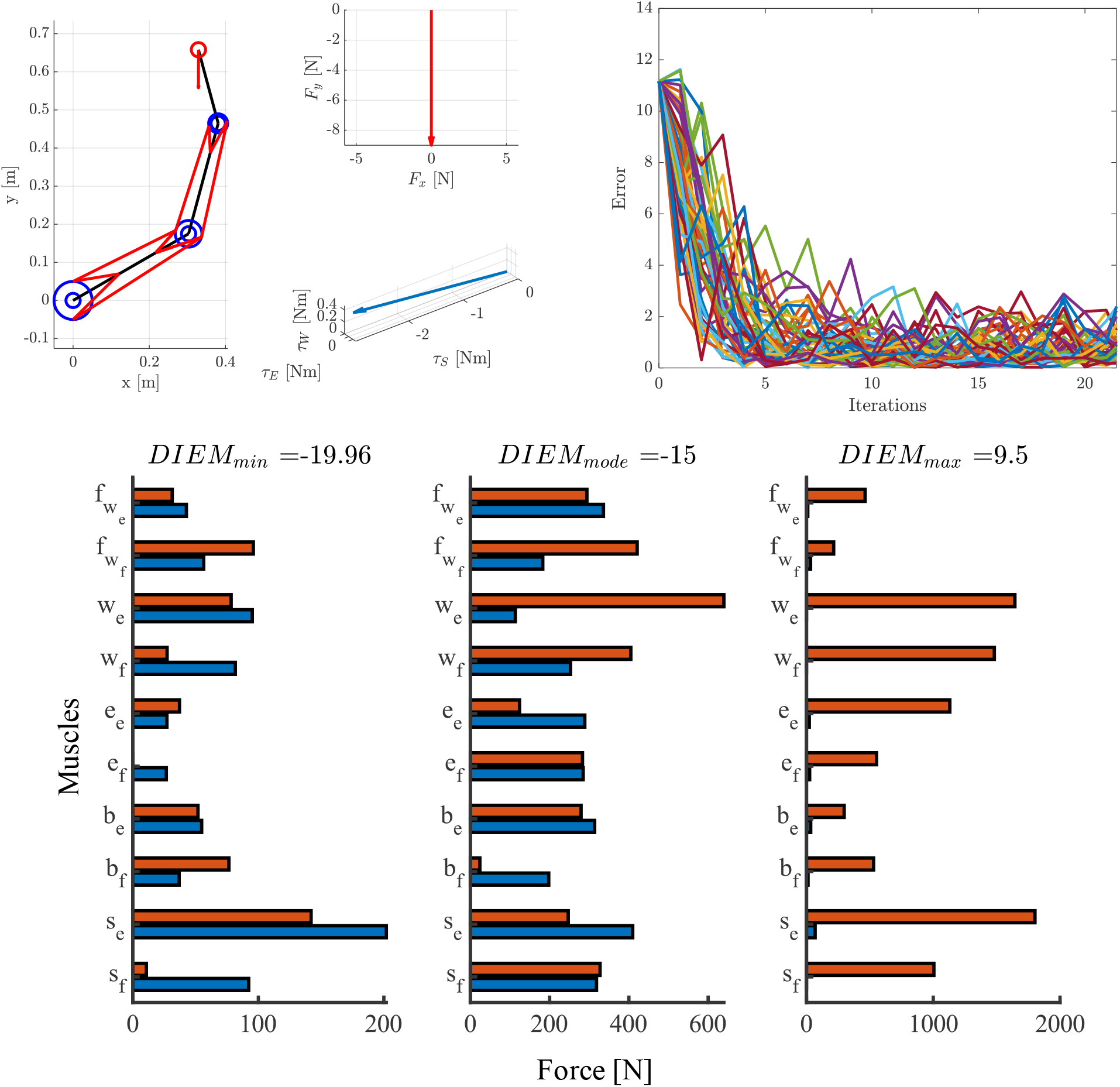
**Top Left**: Three body rigid arm model generating a horizontal arm force. The left panel shows the arm configuration and desired force direction (red arrow). The top right panel show the desir ed end-effector force, while the bottom right panel show the obligatory torque coordination to generate such end -effector force. **Top Right**: Trend of the error function with respect to the number of iterations for the stochastic gradient descent learning algorithm of selectable muscle force synergies. Each colored line represents a different run of the algorithm. **Bottom**: Bar plots presenting the muscles forces of: (left) the two most similar (*DIEM*_*min*_ = −19.96) generated synergies, (center) two synergies near the modal value of the DIEM distribution (*DIEM*_*mean*_ = −15.00) synergies, and (right) the two most dissimilar synergies (*DIEM*_*max*_ = 9.50). These synergies were selected among the 51 different generated synergies. The y-axis labels represent the 10 different muscles: two shoulder mono-articular flexors (*s*_*f*_, *s*_*e*_), two elbow mono-articular flexors (*e*_*f*_, *e*_*e*_), two wrist mono-articular flexors (*w*_*f*_, *w*_*e*_), two bi-articular shoulder-elbow flexors (*b*_*f*_, *b*_*e*_), two mono-articular fore-arm wrist flexors 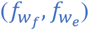.

**Figure 16.**
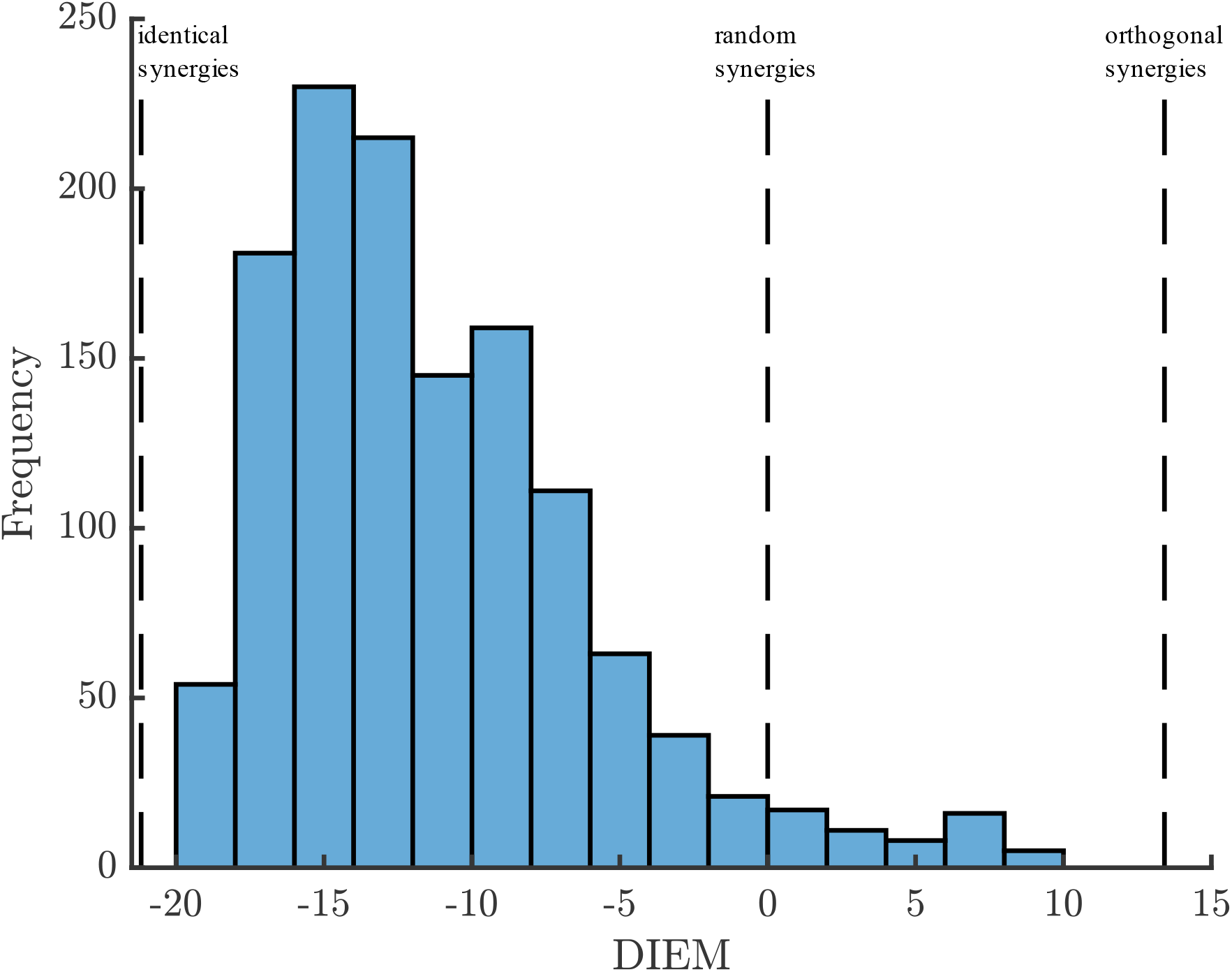
Distribution of the Dimension-Insensitive Euclidean Metric (DIEM) computed between all the 51 different generated muscle synergies. The vertical dashed lines represent the DIEM values respectively for: identical, random, and orthogonal synergies.

### Comparison between Selectable Synergies and the Least Square Solution

For each direction (‘+x,-x,+y,-y’), the 50 selectable synergies were also specifically compared using:

(i) DIEM (hyper-dimensional distance) and (ii) force vector magnitude 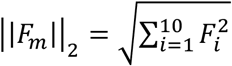.

Figure 17 presents the DIEM histograms between the 50 selectable synergies and the least-square solution. The histograms present similar trends to the ones when all the synergies (51+1) are compared between each other, thus indicating that the selectable synergies did not have a particular relation with the least square solution.

**Figure 17.**
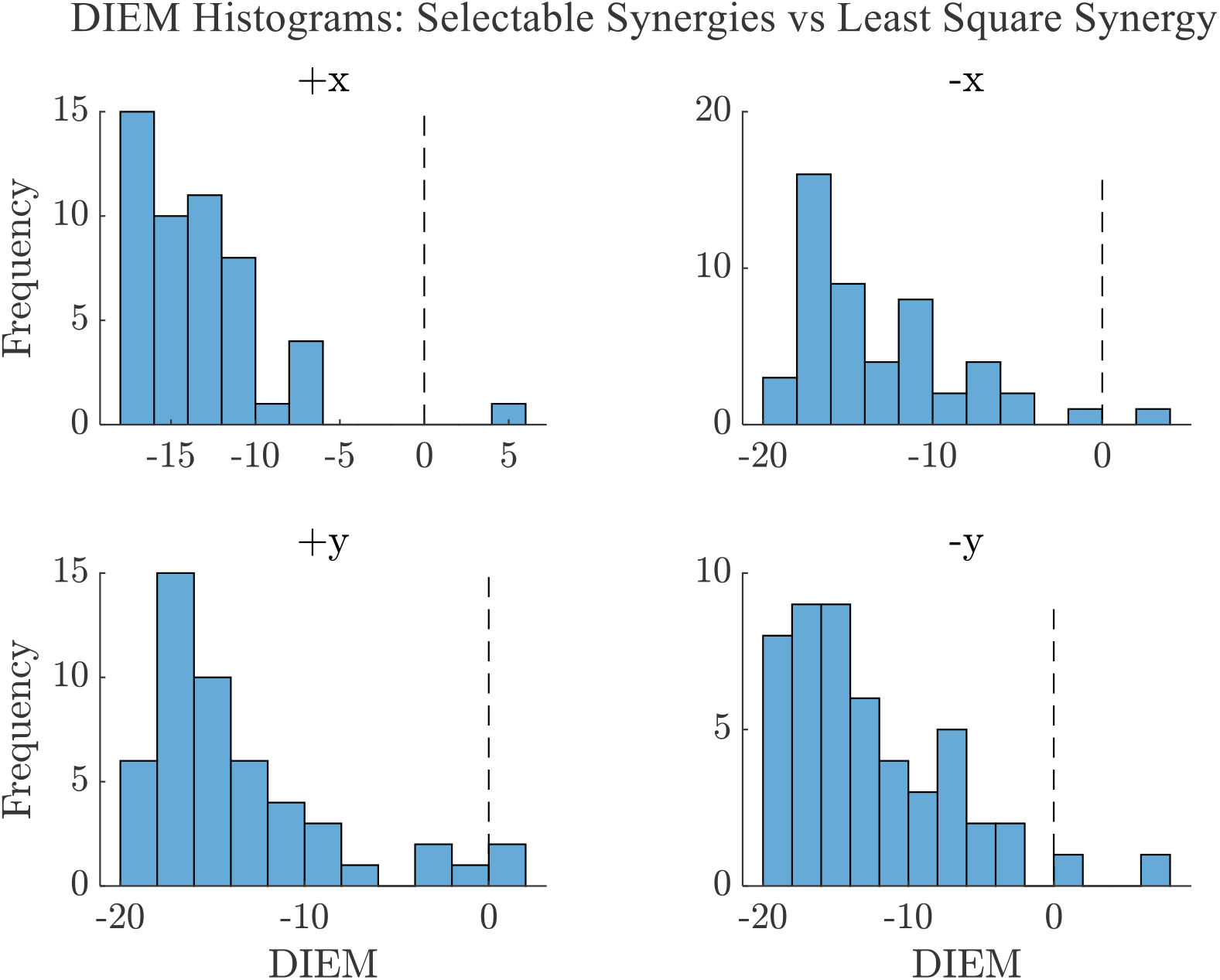
Distribution of the Dimension-Insensitive Euclidean Metric (DIEM) computed between the 50 different selectable muscle synergies and the least square muscle synergy in each of the f our directions: top left +x, top right -x, bottom left +y, bottom right -y. The vertical dashed line represents the DIEM value for random synergies.

Figure 18, instead, shows the force vector magnitude (||*F*_*m*_||_2_) for the 50+1 generated synergies. It can be noted that in most cases, the least square solution has a magnitude lower than all the other synergies. However, this is not always true. Specifically:

**Figure 18.**
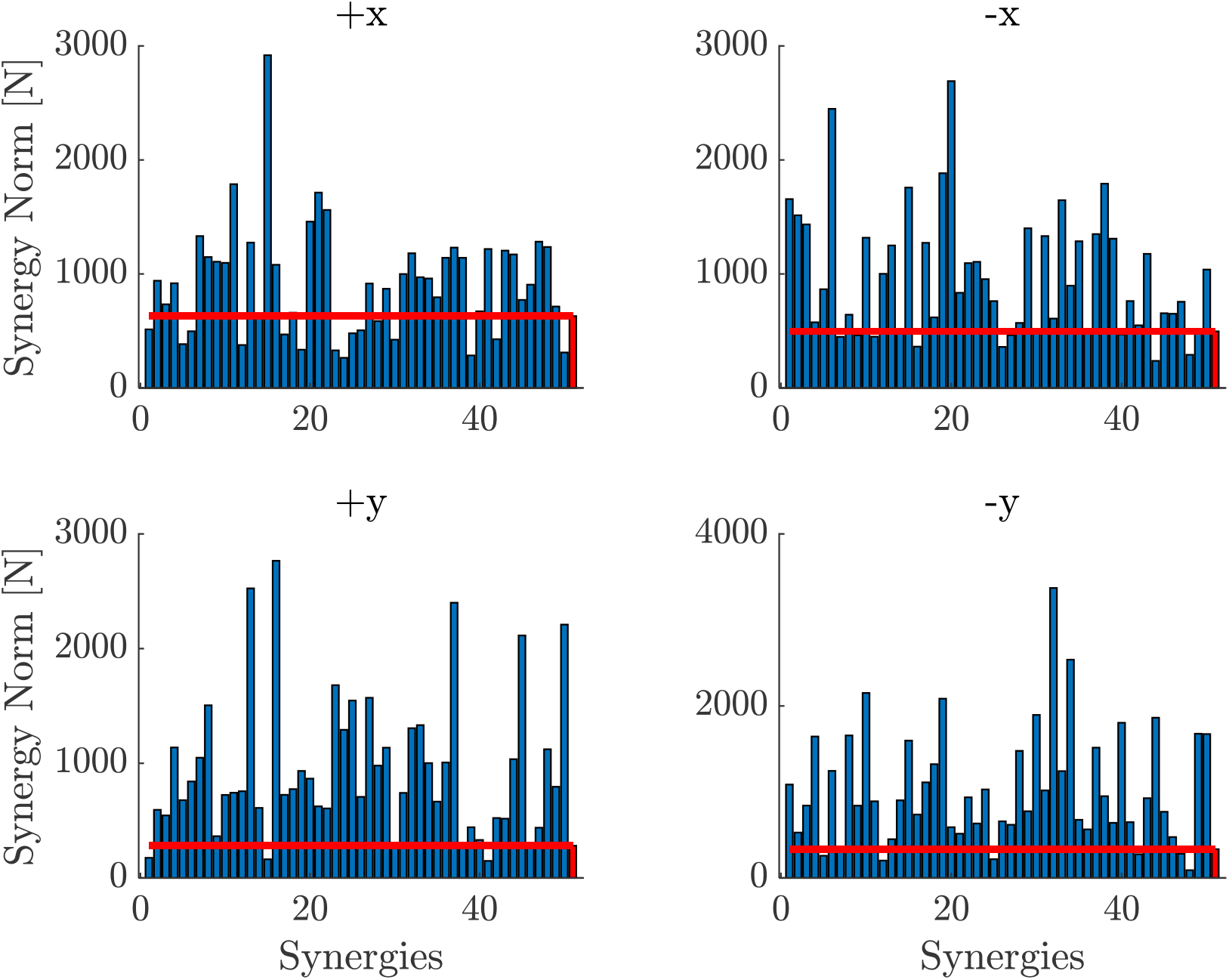
Norm of the synergy vectors (||*F*_*m*_||_2_) in each of the four directions: top left +x, top right -x, bottom left +y, bottom right -y. The first 50 bars (blues) represent the selectable muscle synergies, while the 51^st^ bar (red) shows the least-square solution. The horizontal red line highlights the norm of the least square synergy.

- in the +x direction, 69% of the selectable synergies were larger than the least square synergy;
- in the -x direction, 76% of the selectable synergies were larger than the least square synergy;
- in the +y direction, 92% of the selectable synergies were larger than the least square synergy;
- in the -y direction, 88% of the selectable synergies were larger than the least square synergy.

This might seem counter-intuitive, since the least-square solution should be the solution with minimum norm. However, it is important to stress that the minimum-norm solution – obtained through the Moore-Penrose inverse – achieved an error several orders of magnitudes smaller than the synergies obtained by stochastic gradient descent – which was set to *e*_*th*_ = 1 · 10^−2^*N*^2^*m*^2^ (please refer to the Numerical Example Section in the main document). The selectable synergies occasionally achieved smaller norm by allowing greater error. Figure 19 provides a graphical representation of the final force error in the four different directions.

**Figure 19.**
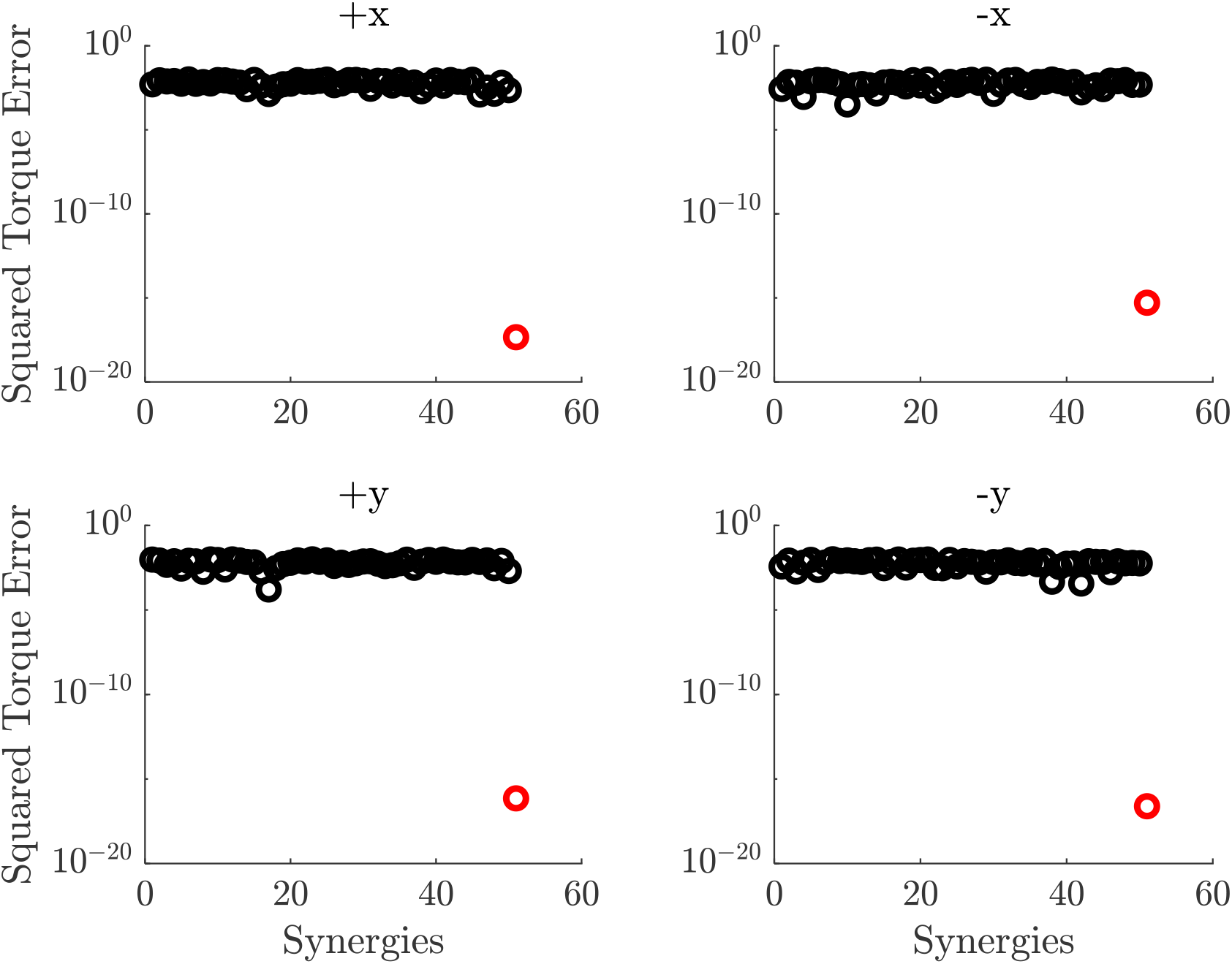
Square torque error computed as expressed in Equation 16 for the 50 selectable muscles synergies (black circles) and for the least square solution (red circle).

## Footnotes

1 One of the most widely adopted training methods for neural network models.

